# Single-nucleus RNA-seq and FISH reveal coordinated transcriptional activity in mammalian myofibers

**DOI:** 10.1101/2020.04.16.043620

**Authors:** Matthieu Dos Santos, Stéphanie Backer, Benjamin Saintpierre, Frederic Relaix, Athanassia Sotiropoulos, Pascal Maire

## Abstract

Skeletal muscle fibers are large syncytia but it is currently unknown whether gene expression is coordinately regulated in their numerous nuclei. By snRNA-seq and snATAC-seq, we showed that slow, fast, myotendinous and neuromuscular junction myonuclei each have different transcriptional programs, associated with distinct chromatin states and combinations of transcription factors. In adult mice, identified myofiber types predominantly express either a slow or one of the three fast isoforms of Myosin heavy chain (MYH) proteins, while a small number of hybrid fibers can express more than one MYH. By snRNA-seq and FISH, we showed that the majority of myonuclei within a myofiber are synchronized, coordinately expressing only one fast *Myh* isoform with a preferential panel of muscle-specific genes. Importantly, this coordination of expression occurs early during post-natal development and depends on innervation. These findings highlight a unique mechanism of coordination of gene expression in a syncytium.

## Introduction

Skeletal muscles constitute 40 to 50% of our body mass and are principally composed of very long myofibers attached to bones via tendons, innervated by motoneurons, and surrounded by vessels and fibroblasts cells. The myofibers themselves are syncytia composed of hundreds of post-mitotic nuclei sharing the same cytoplasm, generated by the fusion of myoblasts during development (Bruusgaard et al., 2003). However, the protein content of myofibers is not homogenous along their length. The proteins required for the formation of the neuromuscular junction (NMJ) accumulate in the center of the myofiber whereas specific proteins for myotendinous junction specialization (MTJ) accumulates at the periphery of the fiber. Two mechanisms could account for this regionalization: the transport of specific proteins to these sites (Kim and Burden, 2008), and the specialization of transcription within the nuclei in these sites (Fontaine et al., 1988). At the NMJ, specific myonuclei express *AchR* and *AchE* (Jacobson et al., 2001) and at the MTJ specific myonuclei express *Col22a1* (Charvet et al., 2013; Koch et al., 2004). However, the precise genetic program operating in these specific myonuclei and the transcription factors involved in this specialization during development, compared to the other myonuclei (called body myonuclei), is far from being understood.

In body myonuclei that constitute the vast majority of myonuclei in the fiber, heterogeneity in nuclear protein import (Cutler et al., 2018) and stochastic gene expression (Newlands et al., 1998) have been proposed to occur, suggesting that body myonuclei along the myofiber are not equivalent. Accordingly, gene transcription could occur stochastically (bursts of transcription), resulting in apparent uncoordinated gene expression at a given locus between the nuclei of the same fiber. Furthermore, in other syncytia such as osteoclasts and syncytiotrophoblasts, transcriptional activity is heterogeneous, varying from one nucleus to the other, indicating that their nuclei, although they share the same cytoplasm, are not coordinated (Ellery et al., 2009; Youn et al., 2010). Whether the hundreds of body nuclei along the myofiber are transcriptionally active at the same time and express the same set of genes remains debated. Adult skeletal muscles are composed mainly of slow and fast myofibers and are usually classified by their MYH (Myosin Heavy Chain) expression profile (Braun and Gautel, 2011; Greising et al., 2012; Gundersen, 2011; Schiaffino and Reggiani, 2011). MYH is the primary determinant of the efficiency of muscle contraction: slow-myofibers in mice express *Myh7* (*MyHC-I*), and fast-myofibers express *Myh2* (*MyHC-IIA*), *Myh1* (*MyHC-IIX*) or *Myh4* (*MyHC-IIB*). These different *Myh* isoforms are coded by different genes, and in mammals, fast *Myh* (f*Myh*) genes including *Myh2, Myh1, Myh4*, embryonic *Myh3*, neonatal *Myh8*, and extraocular *Myh13* are clustered on a single locus while the slow *Myh7* gene lies on another locus (Shrager et al., 2000). Myofibers can express one, two or three MYH isoforms at the same time at protein and mRNA level (Bottinelli et al., 1994; LaFramboise et al., 1991; Murgia et al., 2017). Myofibers expressing more than one MYH isoform are called hybrid fibers in contrast to pure fibers expressing only one isoform. Post-transcriptional mechanisms involving antisense RNA (Rinaldi et al., 2008) or mir-RNA(van Rooij et al., 2009) may participate in the control of the accumulation of specific MYH proteins in both homogenous and hybrid myofibers. Alternatively, hybrid fibers may be constituted of regionalized *Myh* genes expression or of uniform *Myh* genes co-expression along the fiber. For the well-studied *β-Globin* locus whose organization is reminiscent of the f*Myh* locus, it was shown that the LCR establishes contacts and activates a single gene at the locus at a given time (Palstra et al., 2008). It was more recently demonstrated that two adjacent genes can be expressed in the same nucleus at a given time by a shared enhancer (Fukaya et al., 2016). Whether one myonucleus can activate one, two or three f*Myh* genes at the same time remains to be established.

In this report, we sequenced the RNAs and the chromatin accessibility from single nucleus (snRNA-seq and snATAC-seq) isolated from adult skeletal muscles to characterize the transcription profile of known cell populations present in adult skeletal muscles, including FAPS, tenocytes, myogenic stem cells and myonuclei, establishing a blueprint of the transcriptional activity of all its nuclei. We further identified 3 main populations of myonuclei: NMJ, MTJ and body myonuclei with specific transcriptional and chromatin accessibility. Our results showed that most myonuclei express only one f*Myh* gene at the same time. By visualizing *in vivo* in mechanically isolated myofibers the localization of f*Myh* pre-mRNAs, we validated these results showing a coordination of f*Myh* expression in the myonuclei of the majority of myofibers of the fast EDL. We also revealed a minority of hybrid myofibers with nuclei expressing several f*Myh* isoforms at the same time. Those hybrid myofibers were more abundant in slow soleus muscle and during denervation in the fast EDL. Last, we characterized how adult f*Myh* genes become activated during post-natal development and showed the importance of innervation for their coordination.

## Results

### SnRNA-seq identifies several cell populations in adult skeletal muscles

To characterize the transcriptional profile of myonuclei within myofibers, we developed single nucleus RNA-seq (snRNA-seq) experiments using a droplet-based platform on purified nuclei from different skeletal muscles of adult mice (Figures 1A and S1A-D). The origin of the nuclei was identified by specific markers (Figures 1B, S1E and S2A), with 68% of nuclei originating from myofibers, 19% from mesenchymal progenitor cells (or FAPS), and around 13% from other cell types including tenocytes, endothelial and lymphoid cells, smooth muscle cells and myogenic stem cells (Figure S2B). Several populations of myonuclei expressed the muscle-specific *Titin* (*Ttn*) gene, among which MTJ myonuclei (expressing *Col22a1*), NMJ myonuclei (expressing *Ache*), a yet unknown myonuclei subpopulation (expressing *Myh9, Flnc, Runx1*), and body myonuclei accounting for 94% of all myonuclei (Figures 1C and 2A). By comparing gene expression in body nuclei and specialized nuclei, we characterized numerous known (*Ache* and *Etv5* for NMJ (Jacobson et al., 2001); *Coll22a1* for MTJ (Koch et al., 2004) and unknown genes specific to these specialized myonuclei (*Etv4* for NMJ, *Maml2* for MTJ) (Figure 2B). Slow myonuclei expressing *Myh7* clustered independently from the fast *Myh1/2/4* myonuclei. The fast myonuclei formed a large cluster that could be subdivided into several subpopulations, all expressing sarcomeric genes like *Ttn* and various *Myh* isoforms (Figures 1C and 2A). *Myh4*+ myonuclei represented 51% of this fast myonuclei population. We also compared the expression of genes in slow and fast nuclei expressing the different types of *Myh* (Figure 1D). Fast *Myh2* and *Myh1* nuclei showed a similar transcription program, contrasting with slow *Myh7* and very fast *Myh4* nuclei each showing a distinct transcriptional program (Figure 1D). Interestingly, three main categories of *Myh4* nuclei were identified, revealing transcriptional heterogeneity among these nuclei (Figures S3A to S3C). We observed higher expression of *Mical2, Pde4d* and *Sorbs2* in the *Myh4B* myonuclei subgroup, of *Taco1, Neat1* and of *Kpna1* in *Myh4A* myonuclei subgroup, and of *Esrrγ, Agbl1* and *Ptpn3* in *Myh4C* myonuclei subgroup. These *Myh4* nuclei sub-populations were distributed differently in distinct muscles, with a higher percentage of the *Myh4C* population in the fast Tibialis compared to the fast Quadriceps and a higher percentage of *Myh4B* nuclei in the Quadriceps as compared with the Tibialis (Figure S3C). From these experiments, we concluded that *Myh4* nuclei can have different transcription profiles, depending on the muscle, what was presently unsuspected.

**Figure. 1.**
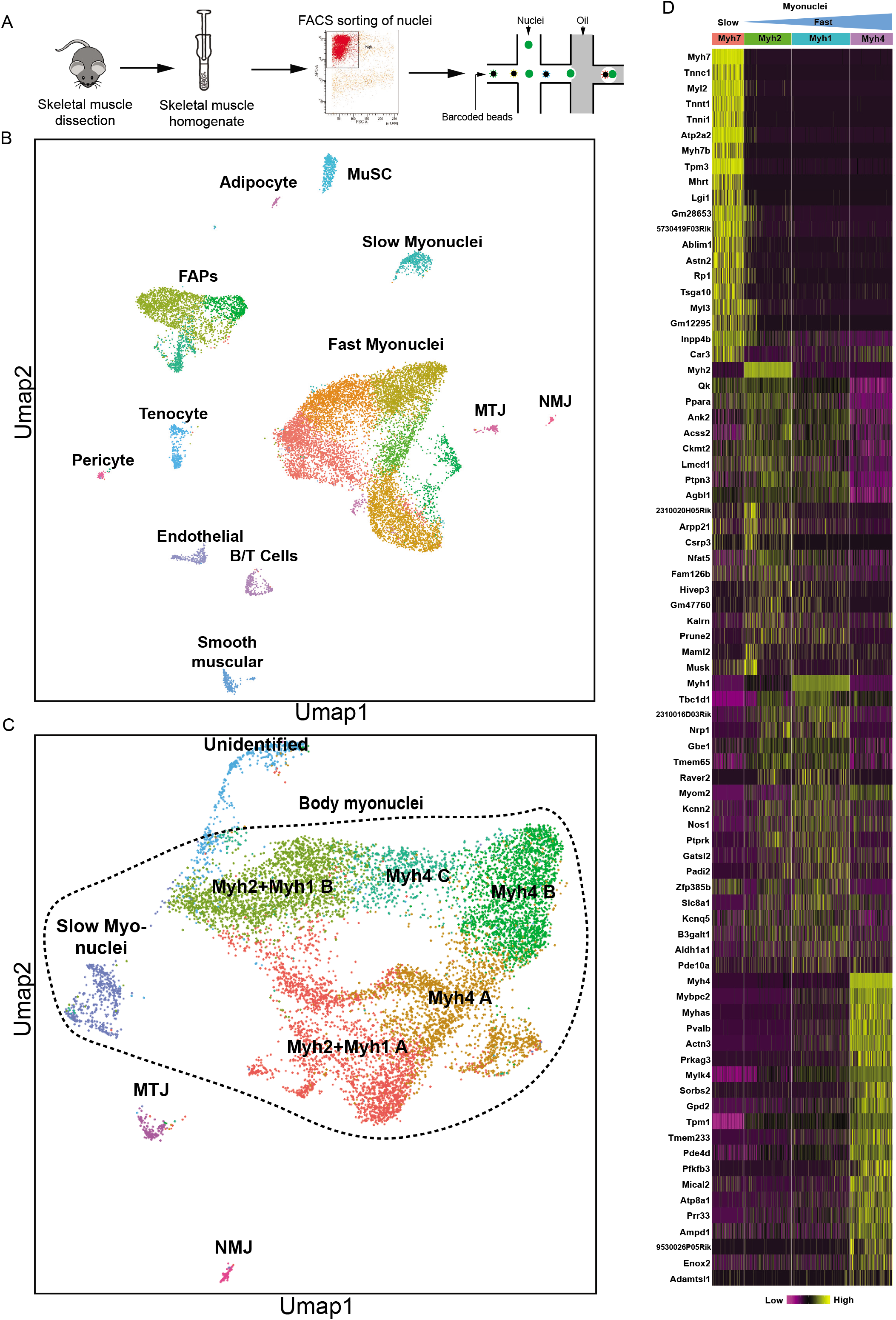
SnRNA-seq analysis from adult slow and fast skeletal muscles. (A) Graphical scheme of the experiments used for snRNA-seq analysis of adult skeletal muscles. (B) Uniform Manifold Approximation and Projection (Umap) diagram from snRNA-seq from adult skeletal muscle. MuSC: Skeletal muscle stem cells; FAPs: Fibro-adipogenic progenitors; MTJ: myotendinous junction; NMJ: neuromuscular junction. (C) Umap diagram of snRNA-seq data with myonuclei only. Myonuclei are separated into 5 clusters: slow, fast, MTJ, NMJ, and unidentified myonuclei. The slow and fast myonuclei are called body nuclei and are surrounded by dotted lines. Five sub-categories of fast myonuclei are clustered. (D) Heatmap of genes upregulated (yellow) and downregulated (violet) in the myonuclei expressing the different slow and fast *Myh* isoforms.

**Figure. 2.**
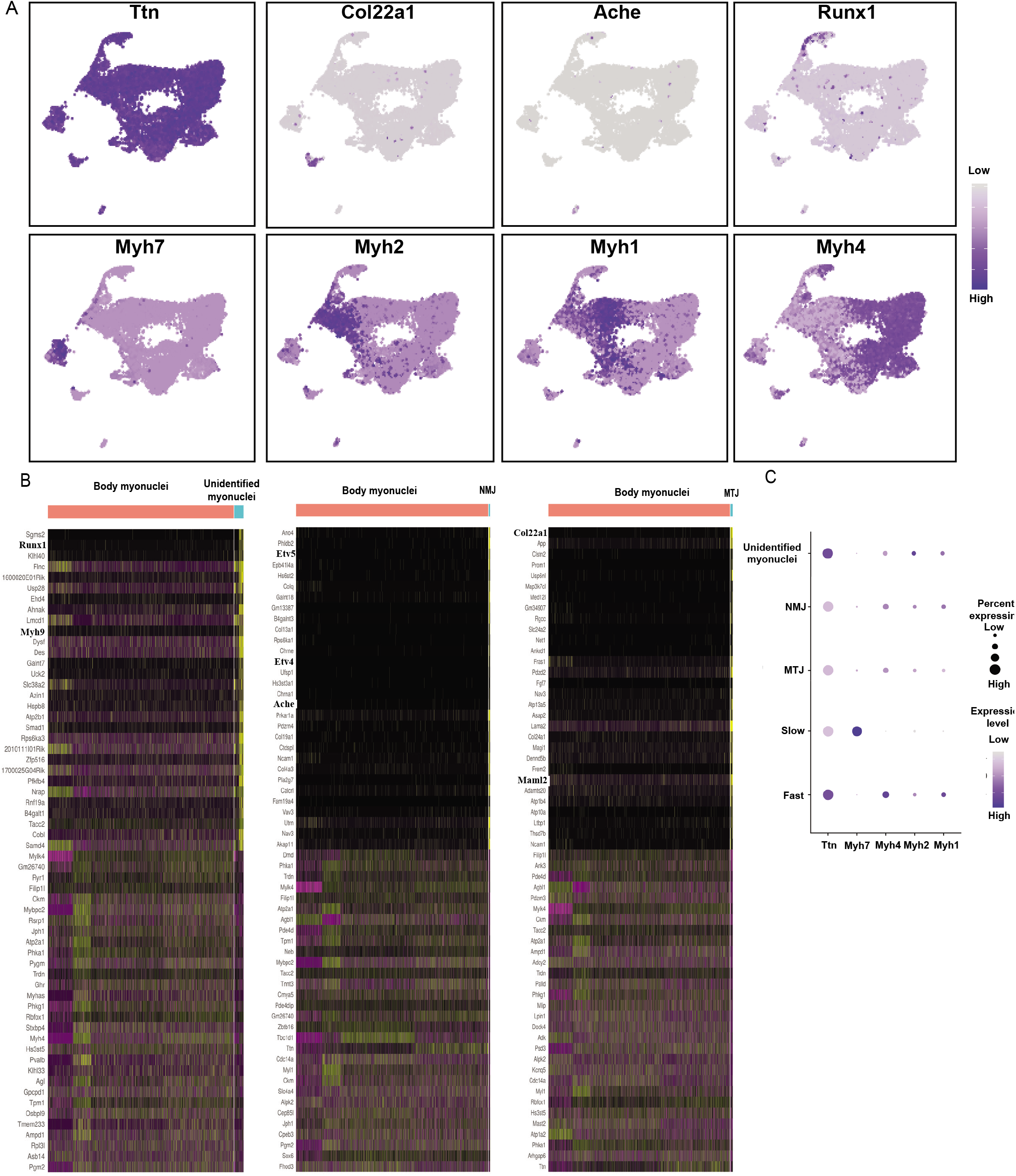
The different populations of myonuclei. (A) Umap Plots showing the expression of several markers used to identify the different types of myonuclei. The intensity of the blue color reflects the level of expression of the gene. (B) Heatmap of the top 30 genes upregulated and downregulated (yellow) in body myonuclei versus unidentified myonuclei (left), in body myonuclei versus NMJ nuclei (center), and body myonuclei versus NMJ nuclei (right). (C) Dot-plots from snRNA-seq experiments showing the expression of the slow and fast *Myh* in the different populations of myonuclei. NMJ and MTJ nuclei expressed *Myh* like other myonuclei.

### SnATAC-seq reveals heterogenous transcriptional programs in myonuclei

Recent studies profiled the genome-wide chromatin accessibility of adult skeletal muscles (Ramachandran et al., 2019). However, this global approach did not discriminate the chromatin status of each cell population present in skeletal muscles. Thus, we set up a protocol to perform single nucleus ATAC-seq (snATAC-seq) with purified nuclei from adult soleus and quadriceps using a droplet-based platform. Briefly, nuclei were transposed using Tn5 transposase and then encapsulated into gel beads emulsion using 10X protocol. After quality assessments and filtering (Figures S4A to S4C), 132 966 peaks were identified from 6037 sequenced nuclei. To identify the different populations of nuclei, we classified nuclei from ATAC-seq based on snRNA-seq experiments from Figure 1B. We used methods for cross-modality integration and label transfer from Signac software (Figure 3A). This classification was robust, as shown by the chromatin accessibility of the *Myh* genes present only in myonuclei (Figures 3B and S4D-E). Using chromVar we characterized the transcription factors (TF) variations in motif accessibility between the different types of myonuclei (Figure 3C). Knowing that this analysis frequently does not distinguish between related TFs of the same family (usually sharing similar motifs) (Schep et al., 2017), we could characterize that the NFAT and SOX binding motifs accessibility are enriched in slow *Myh7* myonuclei, ESRRβ and NR4A2 motifs in fast *Myh2* and *Myh1* myonuclei and SIX and NRL/MAF motifs in fast *Myh4* myonuclei (Figures 3C, 3D and S3B). These distinct combinations of TF could participate in the genetic code controlling the expression of specific fast-subtypes and slow sarcomeric genes and of genes associated with myofiber specialization (Figure 1D). Equivalently to Figure 1D presenting genes differently expressed in slow and fast myonuclei (from snRNA-seq data), we observed specific chromatin accessibility between slow and fast myonuclei in genes differently expressed in slow and fast myonuclei. For exemple, the gene *Pvalb* expressed only in fast *Myh4* myonuclei (Figure 1D) presented a strong chromatin accessibility only in fast *Myh4* myonuclei (Figure S4G). At the opposite a strong chromatin accessibility in the *Atp2a2* was observed in *Myh7* nuclei only where this gene is expressed (Figures 1D and S4H). These results showed for the first time with a good resolution how slow and fast-subtypes genetic programs are associated with different sets of TF controlling the expression and the chromatin accessibility of a set of co-expressed genes. Interestingly, a specific set of TF motifs is enriched in MTJ myonuclei as compared to body myonuclei suggesting their participation in regionalized expression of specific genes such as *Col22a1* (Figures 3E and S4F).

**Figure. 3.**
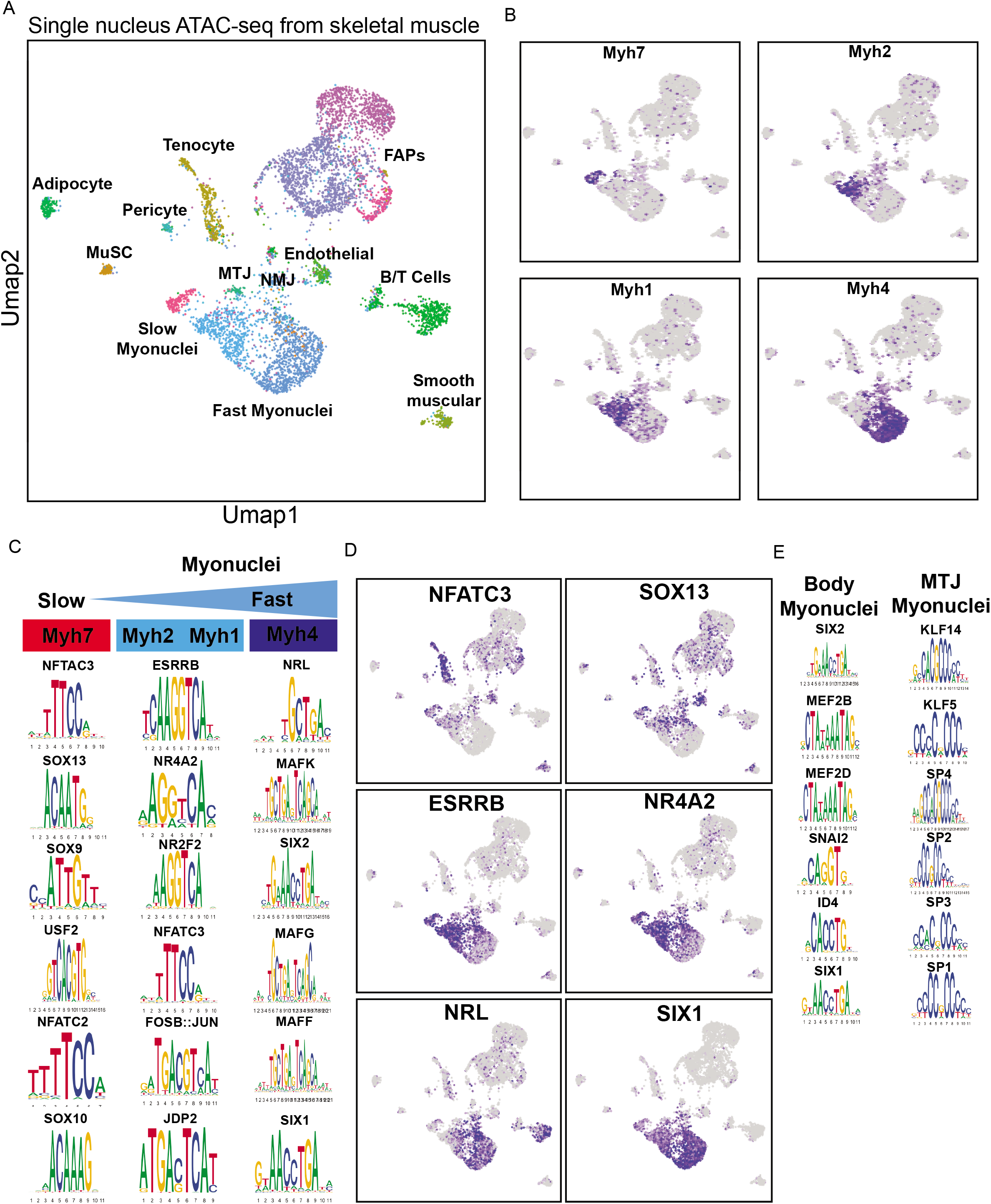
Chromatin accessibility in adult myonuclei by snATAC-seq experiments. (A) Umap Plots from snATAC-seq from adult skeletal muscle based on chromatin accessibility. MuSC: Skeletal muscle stem cells; FAPs: Fibro-adipogenic progenitors; MTJ: myotendinous junction; NMJ: neuromuscular junction. (B) Umap Plots showing the chromatin opening of slow and fast *Myh* genes used to identify the different types of myonuclei. The intensity of the blue color reflects the level of chromatin accessibility of the gene. (C) PWM motifs showing the enrichment of TF motifs in differential peaks in the myonuclei expressing the different *Myh* genes. (D) Umap Plots showing the motif activity of TF in the different types of myonuclei. The intensity of the blue color reflects the level of TF motif enrichment in the nuclei. (E) PWM motif showing the enrichment of TF motifs in differential peaks in the body and MTJ myonuclei.

### Most myonuclei express a single f*Myh* gene

Fast *Myh* genes are located side by side on the same locus and it is presently unknown whether they can be transcribed simultaneously (Figure 4A). Using snRNA-seq, we found that only rare mixed nuclei co-expressed simultaneously *Myh7* and *Myh2* or *Myh1* and *Myh4* (Figure 4B). Mixed nuclei co-expressing *Myh2* and *Myh1* were more numerous. Other combinations of coexpression were not detected (Figure S5A). This showed that the vast majority of adult myonuclei analyzed were pure and expressed only one *Myh* gene. To visualize the expression of fast *Myh* (f*Myh*), we performed fluorescent *in situ* hybridization (RNAscope) experiments using intronic probes to detect f*Myh* pre-mRNA in myofibers from extensor digitorum longus (EDL) and soleus muscles (Figures 4C to 4E). Myonuclei expressed f*Myh* genes in a mono or bi-allelic manner (Figure 4C). Expression of two distinct f*Myh* genes from identical or distinct alleles was detected in some nuclei (Figure 4D). Expression of two f*Myh* genes by the same allele (single bicolor dot) indicated that two *Myh* genes were transcribed simultaneously on the same locus (Figures 4E to 4G). In some myonuclei, no f*Myh* pre-mRNA was detected. Quantification of RNAscope experiments is provided in (Figures 4F to 4G): in EDL, the majority (more than 80%) of nuclei expressed only one *Myh* isoform and less than 5% coexpressed two isoforms. The number of nuclei co-expressing two isoforms of *Myh* was highest in the soleus, where approximately 25% co-expressed *Myh1* and *Myh2* (Figure 4F). The coexpression was in most cases from the same allele (Figure 4G), suggesting that *Myh2* and *Myh1* can be activated simultaneously from this allele. The results of RNAscope experiments thus confirmed snRNA-seq results, showing that almost all myonuclei were pure, expressing only one *Myh* isoform at a time, except in the soleus (Figure 4H). Then we tested whether different populations of myonuclei express either metabolic, calcium handling or sarcomeric genes, or if all myonuclei coexpress simultaneously these different categories of genes. SnRNA-seq data showed that most *Myh4* myonuclei expressed the calcium handling gene *Pvalb* and the glycolytic gene *Pfkfb3*, that a majority of f*Myh* myonuclei expressed *Tbc1d1* involved in muscle glucose uptake, and most *Myh7* nuclei expressed the sarcoplasmic reticulum Calcium transporting 2 gene *Atp2a2* (Figures S6A to S6D). These results were confirmed by RNAscope visualizing the localization of the pre-mRNA of *AldolaseA*, associated with glycolytic metabolism in myofiber (Salminen et al., 1996), and of *Idh2*, associated with oxidative metabolism (Murgia et al., 2015). Accordingly, more than 70% of *Myh4* expressing nuclei also expressed the glycolytic enzyme *AldolaseA* pre-mRNA, while the TCA cycle isocitrate dehydrogenase *Idh2* pre-mRNA was undetectable. At the opposite most *Myh2* expressing nuclei expressed *Idh2* pre-mRNA (Figures S6E to S6G), while *AldolaseA* pre-mRNA was not detected. Thus, myonuclei co-expressed coordinated specialized sarcomeric, calcium handling and metabolic genes. Altogether these results showed that f*Myh* gene expression, and more generally muscle gene expression within the myofiber syncytium is not random but precisely controlled, each myonucleus showing a precise pattern of coexpressed genes.

**Figure. 4.**
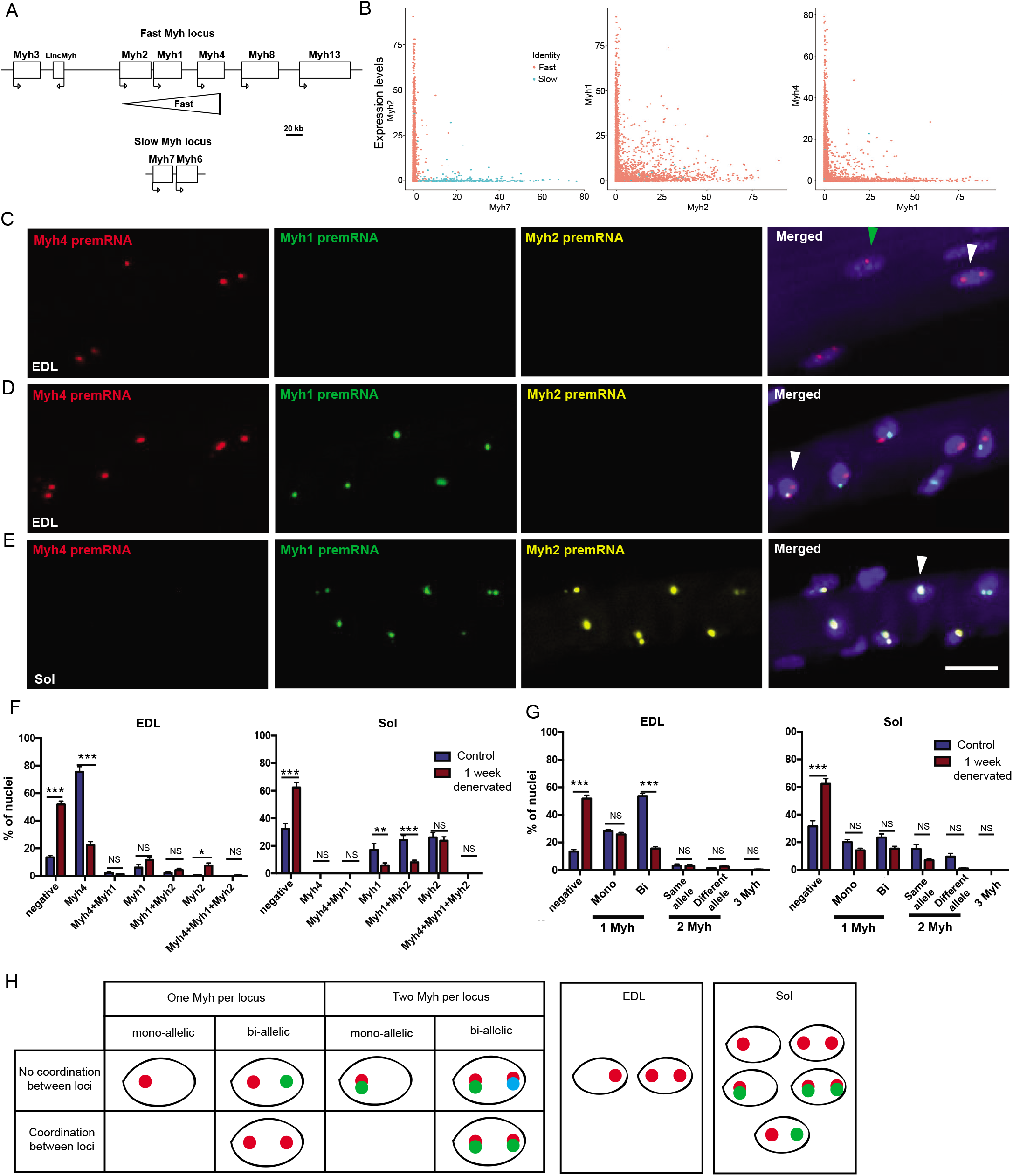
The majority of myonuclei express only one isoform of *Myh*. (A) Diagram of the mouse *Myh* fast and slow loci. (B) Analysis of *Myh* isoforms expression in myonuclei from snRNA-seq data. Each dot corresponds to a myonucleus and the x- and y-axis corresponds to the indicated *Myh* expression level. (C) RNAscope experiments on isolated fibers from EDL to visualize fast *Myh4* (red), *Myh1* (green) and *Myh2* (yellow) pre-mRNAs. *Myh* pre-mRNA can be detected as two transcribed alleles (white arrowhead) or as a single allele (green arrowhead). (D) Same as (C) showing nuclei expressing at the same time different isoforms of *Myh* from each *Myh* allele and nuclei co-expressing (arrowhead) at the same time 2 isoforms of *Myh* from the same allele. (E) Same as (C) in soleus showing nuclei co-expressing at the same time *Myh1* and *Myh2* pre-mRNAs from each allele (arrowhead). (F) Percentage of nuclei expressing f*Myh* pre-mRNAs, in control and one week after denervation, in fast myofibers from EDL and soleus. (G) Percentage of nuclei expressing one, two and three isoforms of pre-mRNAs of *Myh* in control or one week denervated EDL and soleus. Percentage of mono- and bi-allelic expression is also indicated. (H) Diagram showing the different possibilities of *Myh* isoforms expression in a myonucleus and the results of our experiments in EDL and soleus. For C, D and E scale bar: 20μm. For F and G, n=3 and twenty fibers per animal. *P < 0.05, **P < 0.01, ***P < 0.001.

### Transcription is coordinated in myonuclei along the myofiber

We next evaluated the coordination of f*Myh* genes expression in distinct myonuclei of the same fiber. In EDL, coordinated *Myh* expression was detected in all myonuclei in a majority of fibers (Figure 5A, upper panel). In contrast, a large number of uncoordinated fibers was detected in soleus (Figure 5B). Different nuclei within the same soleus fiber were detected expressing different f*Myh* isoforms, while some nuclei co-expressed two isoforms. Quantification of these experiments is provided in (Figures 5C and 5D). We never observed hybrid coordinated myofibers. In quadriceps and EDL, most myofibers (90%) showed *Myh* gene expression coordination between their myonuclei expressing only *Myh4* (Figures 5C and 5D). *Myh4* mRNA accumulated homogenously in the coordinated fibers expressing *Myh4* pre-mRNA (Figure S7A). Interestingly, half of the fibers from the soleus were hybrid (Figure 5C and 5D). In hybrid *Myh4* and *Myh1* fibers, *Myh4* mRNA accumulated less efficiently, and did not spread far from the nuclei expressing *Myh4* pre-mRNA (Figure S7A), demonstrating that mRNA diffusion was confined within myonuclear domains (Cutler et al., 2018). The transcriptional program of *Myh4* positive myonuclei may differ in hybrid myofibers from pure myofibers, explaining the heterogeneity of *Myh4* myonuclei observed by snRNA-seq (Figures S3B and S3C). *Myh4* pre-mRNA was detected also in NMJ myonuclei, as revealed by *AchE* mRNA expression (Figure S7B). Figure 5E summarizes these results.

**Figure. 5.**
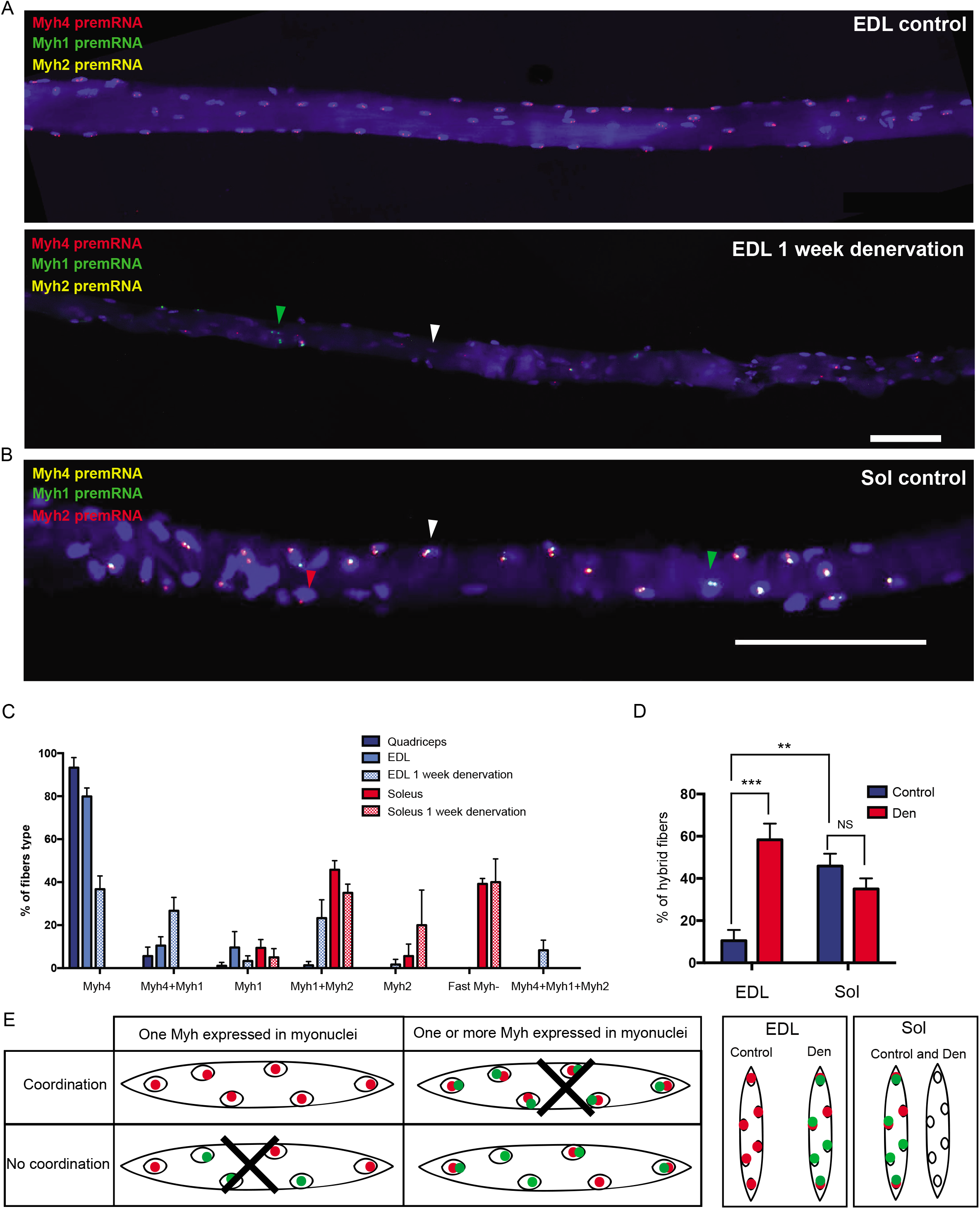
Coordination of fast *Myh* expression along myofibers. (A) RNAscope on isolated fibers from EDL showing the localization of *Myh4* (red), *Myh1* (green) and *Myh2* (yellow) pre-mRNAs. Up: EDL myofiber showing coordination of *Myh4* expression between nuclei (pure myofiber). Down: EDL myofiber after 1 week of denervation. After denervation, myonuclei are no more coordinated in the myofiber and express either *Myh4* or *Myh1* (green arrowhead) or co-express both isoforms (hybrid myofiber). In most myonuclei f*Myh* pre-mRNAs are no more detected (white arrowhead). (B) Same as (A) with *Myh4* (yellow), *Myh1* (green) and *Myh2* (red) pre-mRNAs in a soleus hybrid myofiber. The white arrowhead shows a mixed *Myh1, Myh2* myonucleus whereas the red arrowhead shows a pure *Myh2* myonucleus. (C) Percentage of the different types of fibers observed in quadriceps, EDL and soleus, and one week after denervation. (D) Percentage of hybrid (de-coordinated) fibers in control EDL and soleus and after one week of denervation. (E) Diagram showing the different possibilities of *Myh* isoforms expression patterns in a myofiber and the results of our experiments in control and denervated EDL and soleus. Black crosses indicate expression patterns not observed. For A-B-C scale bar: 100μm. For D-E, n=3 and twenty fibers per animal. *P < 0.05, **P < 0.01, ***P < 0.001.

### Innervation controls myonuclei coordination along the myofiber

To understand the basis behind pure and mixed myonuclei and their coordination along the myofiber, we tested whether innervation, known to modulate adult muscle fiber type (Gundersen, 2011; Rowan et al., 2012; Schiaffino and Reggiani, 2011), was involved in the coordination process. At the protein level, one week after denervation, we observed a preferential atrophy of fast MYH4 and MYH1 positive fibers and 4 weeks after denervation a fast to slow transition in all muscles (Figure S8A). In agreement, one week after denervation, the number of nuclei negative for fast *Myh* genes was increased (Figures 4F and 5A). The percentage of uncoordinated myofibers after one-week denervation in the EDL was increased more than five-fold but did not significantly differ in soleus (Figure 5D). These observations showed that innervation is absolutely required to coordinate fast *Myh* genes expression in the nuclei within myofibers in EDL (Figures 5A to 5D, and S8D). Shown in Figure S8E, a strong increase of *Myh2* and *Myh1* pre-mRNAs was detected in the MTJ myonuclei of EDL one week after denervation, suggesting that in absence of innervation or muscle contraction, MTJ myonuclei expressed a distinct *Myh* gene than body myonuclei (Dix and Eisenberg, 1990). These data support an important role of innervation in the activation of f*Myh* genes transcription and coordination in myonuclei along the myofiber. Altogether these experiments revealed that in normal conditions myonuclei of fast myofibers (Quadriceps and EDL) are coordinated and that denervation induced an increased number of de-coordinated hybrid fibers associated with a shift of *Myh* expression toward slower isoform. In the soleus more hybrid fibers are present and denervation did not induce drastic changes of f*Myh* expression like the ones observed in fast muscles. This suggests that the expression of *Myh1* and *Myh2* in the soleus is less under the control of innervation than the expression of *Myh4* in fast muscles. Knowing that in adult myofibers the homeoprotein SIX1 controls the expression of many fast/glycolytic muscle genes among which f*Myh* genes (Sakakibara et al., 2014; Sakakibara et al., 2016) and that SIX binding motifs accessibility are enriched in *Myh4* myonuclei ((Ramachandran et al., 2019), Figure 3C), we looked at the nuclear accumulation of this protein in fast myonuclei along EDL myofibers. As shown in Figure S9A we observed robust SIX1 accumulation in all myonuclei from an extremity to the other of the fiber. To further test the involvement of SIX1 in f*Myh* gene coordination along the myofiber we tested the consequences of its absence in mutant myofibers from muscle specific *Six1* conditional knock-out (Figures S9B and S9C). In absence of *Six1*, EDL myofibers appeared slower and presented an increased number of non-coordinated hybrids fibers (Figures S9B to S9C), showing that SIX transcription complexes participate to coordinate the expression of f*Myh* genes in adult myonuclei.

### Coordination of myonuclei during development and adult regeneration

We next addressed how f*Myh* expression coordination appears during development. We analyzed the expression pattern of mRNAs or pre-mRNAs of all sarcomeric *Myh* (embryonic, neonatal, slow and adult fast) on isolated forelimbs myofibers at different stages. At E15.5, when myofibers were already innervated, embryonic primary myofibers expressed *Myh3* and *Myh8* mRNA (Figures S10A and S10B). At E18.5, secondary fetal myofibers (small fibers) displayed a higher accumulation of *Myh3* than of *Myh8. Myh3* mRNAs accumulated strongly in MTJ regions of myofibers at all stages, (Figures S10A and S10B) and after birth also at the center of myofibers, where NMJs are suspected to be localized (Figure 6A and S10A). These results suggested the specific transient activation of *Myh3* in NMJ and MTJ myonuclei, or its transient activation in newly accreted myonuclei in these regions of growing myofibers as proposed previously (Zhang and McLennan, 1995). In contrast, *Myh8* mRNA was not regionalized along the myofiber at any stage. Adult f*Myh* pre-mRNAs were first detected in a few nuclei at E15.5 and their number increased drastically after E18.5 (Figures 6B and 6C). At these different stages, most of the myonuclei expressed only one adult f*Myh* gene (Figure 6D) without obvious regionalization along the myofiber. We observed mono-allelic expression before E18.5, and bi-allelic expression after P2 (higher transcription rates) (Figure 6E). At P2 and P5, more than 90% of the nuclei were coordinated along the myofibers and expressed one f*Myh* gene (Figure 6F). This percentage was similar to that of adult EDL, suggesting that f*Myh* gene coordination is established early when the expression is initiated. *Myh2* and *Myh4* mRNAs accumulated strongly from P5 but some areas of the fiber did not present mRNAs accumulation (Figure S11A). Slow *Myh7* mRNAs accumulated in all primary fibers at the MTJ level at E15.5 (Figure 6G). After E18.5, the expression of *Myh7* was restricted to slow fibers which homogeneously accumulated this mRNA specifically (Figure S11B). These experiments showed that the majority of myonuclei started to express adult f*Myh* genes at P2, a single gene being activated by nucleus, and that myonuclei in the growing myofiber were coordinated from their onset. We next wonder how adult f*Myh* were expressed during muscle regeneration (following a lesion), in which MuSCs rebuild newly formed myofibers. Strikingly, at the opposite of what we observed during development, 7 days after adult muscle injury, most of the regenerating myofibers were hybrid fibers with nuclei expressing different isoforms of f*Myh* (Figure 6H). Remarkably these regenerating fibers showed a regionalization of nuclei expressing distinct f*Myh* genes, suggesting either the involvement of heterogenous signaling from surrounding cells along the myofiber or the fusion of distinct preprogrammed myogenic stem cells having fused together and not yet re-coordinated by motoneuron firing input (Kalhovde et al., 2005).

**Figure 6.**
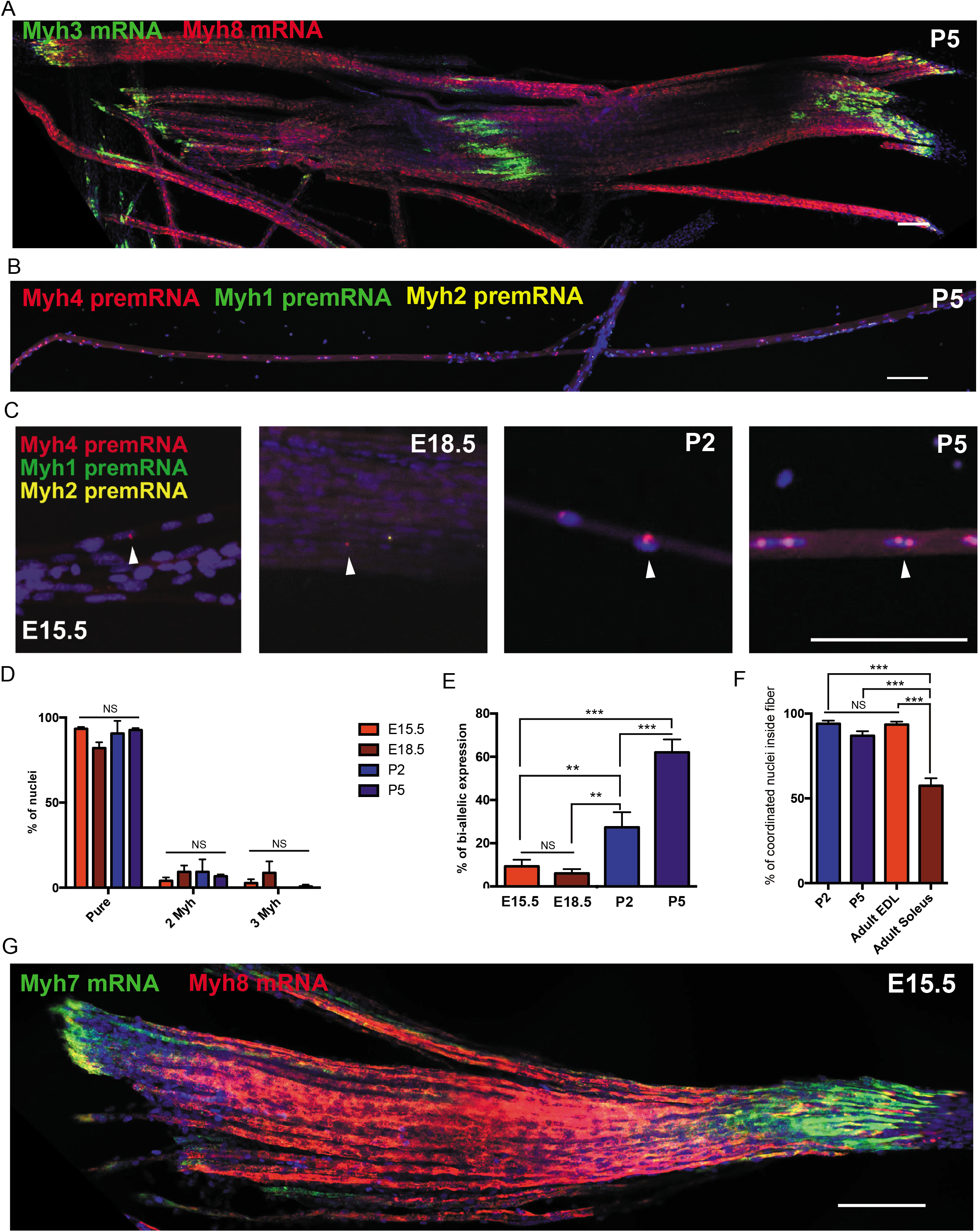
f*Myh* expression during development and regeneration is regionalized. (A) RNAscope against *Myh3* (green) and *Myh8* (red) mRNA on isolated fibers from forelimbs at 5 days post-natal (P5). The MTJ and NMJ areas showed an accumulation of *Myh3* mRNA. (B) RNAscope against *Myh4* (red), *Myh1* (green) and *Myh2* (yellow) pre-mRNAs on isolated fibers at E15.5, E18.5, 2 (P2) and 5 (P5) days post-natal. (C) Same as (B). Zoom showing the increase of adult fast *Myh* positive (arrowhead) nuclei after birth. (D) Percentage of nuclei expressing one, two or three adult f*Myh* isoforms at the same time at different developmental stages. (E) Percentage of nuclei with bi-allelic expression of *Myh* genes during development. (F) Percentage of coordinated nuclei inside myofibers at different developmental stages. (G) RNAscope against *Myh7* (green) and *Myh8* (red) mRNAs on isolated fibers at E15.5. *Myh7* mRNAs accumulate in MTJ areas of all myofibers at E15.5. (H) RNAscope experiments on isolated fibers from 7 days regenerating EDL to visualize fast *Myh4* (red), *Myh1* (green) and *Myh2* (yellow) pre-mRNAs. For A-B-C-G scale bar: 100 μm. For D, n=3 and 50 nuclei per animal. For E-F, n=3 and twenty fibers per animal. *P < 0.05, **P < 0.01, ***P < 0.001.

## Discussion

By combining single-nucleus RNA-seq/ATAC-seq and FISH experiments, we established for the first time a complete atlas of the gene expression pattern and chromatin landscape of the different cell populations of adult skeletal muscles, including mononucleated cells and multinucleated myofibers. From our experiments, we estimated that myonuclei constitute approximately 68% of the nuclei present in adult skeletal muscles of two months old mice at rest. Recent single-cell RNA-seq studies of skeletal muscle (Dell’Orso et al., 2019; Giordani et al., 2019) have characterized the gene expression pattern of mononucleated cells of adult skeletal muscle, but failed to identify the transcription signature of multinucleated myofibers. The composition and gene expression of cell types identified by these reports is in agreement with our data, providing the same global signature of skeletal muscle at rest, identifying distinct populations of FAPS, tenocytes, immune cells, pericytes adipocytes, smooth muscle cells and myogenic stem cells. Expanding these studies, our results showed that myofibers are composed of three different types of myonuclei localized differently and expressing distinct sets of genes (Figure 2), namely the MTJ, NMJ and body myonuclei. The full transcriptome of these nuclei was to this day unknown, and we characterized numerous genes expressed specifically in these specialized myonuclei. We also characterized an unidentified population of *Titin+* myonuclei expressing *Runx1, Flnc* and *Myh9*. As FilaminC is involved in early stages of myofibrillar remodeling and repair, this myonuclei population which had never been described could correspond to myonuclei recently fused to the myofiber or to myonuclei in area of myofiber damage to support fast repair (Leber et al., 2016). Our results demonstrated that most myonuclei express a single f*Myh* gene, and that in the majority of myofibers myonuclei coordinately express a specific set of genes including this *Myh* gene. Our RNAscope experiments also identified a unique feature of transcriptional control in myonuclei of adult myofibers. In fast skeletal muscles, the transcriptional activity of nuclei within each myofiber is finely coordinated. We showed that this coordination is dependent upon innervation and is established early during development.

Several reports have shown that apparently homogenous populations of cells present an important variability of gene expression. This variability can be the consequence of stochastic gene expression, different stages of cell cycle, or distinct subcategories of cells in the studied population (Martinez-Jimenez et al., 2017). In syncytia like osteoclasts and syncytiotrophoblasts, gene expression can be heterogeneous in nuclei that share the same cytoplasm (Ellery et al., 2009; Youn et al., 2010). We show here that it is not the case for adult myofibers. We found that the transcription of adult f*Myh* genes in myonuclei along given myofibers was finely controlled during development, allowing the coordinated expression in their nuclei of a single f*Myh* gene. This transcriptional coordination could be a consequence of various processes. In *Drosophila*, one founder myonucleus was shown to reprogram newly fused myonuclei, inducing coordination of expression between nuclei of the same fiber (Bataille et al., 2017). Alternatively, distinct populations of preprogrammed myogenic stem cells with distinct potential (Kalhovde et al., 2005; Lee et al., 2015) could fuse (homotypic fusion (Pin et al., 2002)), activating a single *Myh* gene along the myofiber. Moreover, extracellular signals from surrounding cells may also intervene (Anakwe et al., 2003). The de-coordination of myonuclei after denervation observed in our study supports the importance of innervation in the coordination process during development. A better understanding of the matching between motoneuron subtypes and myofiber subtypes that operates during development (Rafuse and Landmesser, 2000) may help understand how neuromuscular specialization contributes to the onset of coordination. Additionally, the reinnervation process during regeneration may contribute to the recoordination of myonuclei along the regenerating myofiber by a community effect (Gurdon, 1988), with all the myonuclei of the myofiber differentiating simultaneously in response to common signaling factors.

Adult myofibers are plastic and can adapt their contractile and metabolic properties depending on external stimuli by activating or repressing a set of specific genes, such as the *Myh* genes (Gundersen, 2011; Schiaffino and Reggiani, 2011). These changes occur in a sequential order recapitulating the organization of the genes at the fast *Myh* locus (*Myh4-Myh1-Myh2*) (Schiaffino and Reggiani, 2011). In this report, we showed that most myonuclei expressed only one *Myh* isoform at a given time (Figure 4). This robust and exclusive expression is reminiscent of the control of the *β-Globin* genes by a locus control region (Palstra et al., 2008). The regulation of the f*Myh* locus is not fully understood and could be controlled like the *β-Globin* genes by a locus control region, such as a super enhancer (Hnisz et al., 2017; Whyte et al., 2013), activating the expression of associated embryonic, fetal and adult *Globin* genes in a temporal order. Transient transfection of adult mouse TA and soleus with *GFP* reporters under the control of 1kb upstream of each *Myh2*, *Myh1* and *Myh4* promoters recapitulate their specific expression in the corresponding MYH2, MYH1 and MYH4 myofibers, respectively (Allen et al., 2001; Allen et al., 2005). This suggests that each MYH promoter possesses its own fiber specific cis-regulatory elements. Accordingly, in *Myh4+* nuclei, we observed in snATAC-seq experiments a higher chromatin opening in *Myh4* than in *Myh1* and *Myh2* genes. In *Myh4+nuclei*, we failed to detect chromatin opening at the *Myh1* and *Myh2* promoter regions, suggesting the absence of TF binding. In soleus myofibers with a high proportion of *Myh1* and *Myh2* coexpression (where *Myh1* and *Myh2* pre-mRNAs are detected in many myonuclei), we mainly observed *Myh1* and *Myh2* pre-mRNAs as two dots in the myonuclei suggesting that each allele transcribed a single gene, and that these two adjacent f*Myh* genes could not be transcribed simultaneously. Whether this exclusive expression reflects the competition for a shared enhancer at the locus, as already illustrated for the *β-Globin* genes (Palstra et al., 2008), is an interesting possibility.

The coordinated expression of a single isoform of *Myh* could be due to a similar combinations of transcription activators and repressors concentration in these myonuclei, affecting all the genes that they express, or only those with high expression, excluding stochastic expression as previously proposed (Newlands et al., 1998). In support of this, snRNA-seq experiments showed that specific genes are co-expressed with *Myh4*, such as genes coding for other sarcomeric proteins, proteins associated with metabolism or associated with calcium handling. The snRNA-seq results showed harmonious non-random gene expression pattern in each nucleus of the myofiber, with some variability between muscles with distinct subprograms among *Myh4+* myonuclei. In the mouse, denervation induces muscle atrophy, associated with a fast to slow transition of *Myh* at the protein level. These changes occur also at the pre-mRNA level, as shown in the present study. We observed that transitions after denervation were sequential, *Myh4+* fibers expressing *Myh1* but not *Myh2*, and some nuclei switching their *Myh* expression profile earlier than others. Thirty days after denervation, myofibers remained uncoordinated showing that the increased number of hybrid fibers observed in EDL 7 days after denervation was not due to a transitional state but that innervation was required to maintain the coordination of f*Myh* expression. The atrophy of myofibers induced by denervation involves the activation of protein degradation pathways and the repression of protein synthesis (Schiaffino et al., 2013). We observed an increased number of f*Myh*-negative nuclei as well as a decreased bi-allelic expression of f*Myh* after muscle denervation, suggesting that down-regulation of f*Myh* expression may participate as well in this atrophy.

The loss of coordination in myonuclei of adult myofibers after denervation observed in the present study is consistent with the role of motoneurons in controlling adult muscle physiology (Gundersen, 2011; Hennig and Lomo, 1985; Rowan et al., 2012; Salviati et al., 1986; Schiaffino and Reggiani, 2011). Calcium is a diffusible signal distributed throughout myofibers and a potential candidate for the control specific muscle gene coordination. Resting Ca^2+^ concentration differs in fast and slow fibers, and is modulated by slow or fast motoneuron firing (Eccles et al., 1957; Eusebi et al., 1980; Olson and Williams, 2000). Ca^2+^ transients following slow motoneuron stimulation activate specific signaling pathways (Gundersen, 2011; Wu et al., 2002) among which Calcineurin phosphatase that controls the nucleo-cytoplasmic localization of NFATc1 in slow myofibers important for the expression of slow muscle genes (Chin et al., 1998; Tothova et al., 2006). MYOD activity as well is modulated by motoneuron activity and phosphorylation of its T115 by slow motoneuron activation impairs its DNA binding activity in the soleus (Ekmark et al., 2007). In contrast, fast-twitch motoneurons, characterized by phasic high frequency firing do not activate Calcineurin phosphatase, suggesting that other signaling pathways relay motoneurons firing information in fast myofibers. Furthermore, fast subtypes myofibers are innervated by fast subtypes motoneurons, including fast fatigable (FF, innervating MYH4 myofibers), fast intermediate (FI, innervating MYH1 myofibers) and fast resistant (FR, innervating MYH2 myofibers) (Fogarty et al., 2019; Kanning et al., 2010; Muller et al., 2014). How these distinct classes of fast motoneurons control the fast-muscle subtypes phenotype and myonuclei coordination is presently unknown (Kanning et al., 2010; Pun et al., 2006; Salviati et al., 1986). Whether Ca^2+^ concentrations are similar along the myofiber and how denervation may perturb this basal intracellular Ca^2+^ concentrations are potential important issues to understand myonuclei coordination. Several proteins controlling intracellular Ca^2+^ concentration in myofibers (Calsequestrin 1 et 2, Serca1 and 2, Parvalbumin, Calmodulin, Phospholamban, Myozenin 1 and 2, Sarcalumenin, IP3R) are differentially expressed in fast and slow myofibers, and induce a variety of local Ca^2+^ concentration gradients that may modulate the activity of different transcription factors (Berchtold et al., 2000; Casas et al., 2010; Frey et al., 2008; Racay et al., 2006). Here, we showed that the transcription factor SIX1 accumulates in all myonuclei of EDL myofibers and participates in the coordination of f*Myh* expression. The mechanisms controlling SIX1 nuclear accumulation specifically in fast myofibers remain to be established. We observed by snRNA-seq an increased expression of the SIX cofactor EYA4 in *Myh4+* myonuclei. Whether the SIX1-EYA4 transcription complex activity may be controlled by motoneuron firing frequency and participates in myonuclei coordination is an interesting possibility that may explain its role in the control of the adult fast phenotype (Maire et al., 2020).

The genome-wide chromatin landscape of adult muscles had not been evaluated previously at the myofiber level, due to the lack of adapted experimental protocols (Ramachandran et al., 2019). We show here that snATAC-seq is a powerful tool to overcome this limitation and that it allows to detect all the cell populations identified by snRNA-seq in adult skeletal muscles, including myofibers. We identified chromatin accessibility domains coherent with the defined patterns of gene expression, specific of fast-subtypes or slow myonuclei, of MTJ or body myonuclei, as well as of other cell types in skeletal muscle. This analysis also allowed to identify enriched TF binding sites in ATAC-peaks specific for each myonuclei population. We identified TF binding motifs known to participate in muscle genes regulation in slow and fast myofibers, among which the NFATc (Tothova et al., 2006), MEF2 (Potthoff and Olson, 2007) and SIX (Grifone et al., 2004; Sakakibara et al., 2014; Santolini et al., 2016) families, and nuclear receptors of the ERR-β/γ (Rangwala et al., 2010) and the NR4A1 (Chao et al., 2007) families. The differential TF ATAC-peaks enrichment observed between fast subtypes myonuclei sheds some light on the basis of their specialization. We identified TF DNA binding motifs which had never been implicated in muscle genes regulation, such as those recognized by the b-ZIP family of NRL/MAF transcription factors (Katsuoka and Yamamoto, 2016). Accordingly, we identified *c-Maf* mRNAs enriched in the *Myh4+* myonuclei in our snRNA-seq, suggesting that this TF may contribute to the fine-tuning of muscle gene expression in specialized fast myonuclei, in synergy with SIX homeoproteins. Serria et al showed earlier that *c-Maf* expression is upregulated in myogenic cells during their differentiation in culture, but the role of *c-Maf* in adult muscle physiology is presently unknown (Serria et al., 2003).

Last, in addition to defining the exact transcriptional landscape within skeletal muscle, the snRNA-seq and snATAC-seq methods presented here open new perspectives to characterize in the future the crosstalk between myofibers and their environment, such as with associated myogenic stem cells in pathophysiological conditions, to date poorly investigated due to technical limits.

## Acknowledgments

We thank P-A. Defossez, E. Bloch-Gallego, P. Mourikis, V.Hakim, D.Duprez and S. Gautron for helpful critical reading of the manuscript. We thank T. Guilbert and F. Letourneur at the Cochin IMAG’IC and GENOM’IC platform for helpful advice. We thank S. Joshi for helpful advice for nuclei purification, and P. Mourikis and L. Machado for helpful discussions. Funding: M.D.S was successively supported by a PhD fellowship from the Association Française contre les Myopathies (AFM), by a post doc fellowship from EUR-Gene and from the Agence Nationale pour la Recherche (Myolinc, ANR R17062KK). Financial support to this work was provided by the AFM (n°17406 to P.M, and 19507 Translamuscle to F.R), the ANR (R17062KK), the Institut National de la Santé et de la Recherche Médicale (INSERM) and the Centre National de la Recherche Scientifique (CNRS).

## Author contributions

M.D.S, A.S, F.R and P.M conceived the study. M.D.S, S.B, B.S and A.S performed experiments. M.D.S and P.M wrote the manuscript with input from A.S and F.R. Competing interests: the authors declare no competing interests. Data and materials availability: all experimental data are available in the main text or the supplementary materials.

## Materials and Methods

### Animals

Animal experimentations were carried out in strict accordance with the European STE 123 and the French national charter on the Ethics of Animal Experimentation. Protocols were approved by the Ethical Committee of Animal Experiments of the Institut Cochin, CNRS UMR 8104, INSERM U1016, and by the Ministère de l’éducation nationale, de l’enseignement et de la recherche, n° APAFIS#15699-2018021516569195. We used 6 to 8 weeks old C57black6N mouse female for most of our experiments. Mice were anesthetized with intraperitoneal injections of ketamine and xylazine and with subcutaneous buprecare injections before denervation was performed by sectioning of the sciatic nerve in one leg. For skeletal muscle regeneration, mice were anesthetized with intraperitoneal injections of ketamine and xylazine and with subcutaneous buprecare injections before injection of 30μl of cardiotoxin 12μM (Latoxan).

### FACS sorting of nuclei from adult skeletal muscle for single-nucleus RNA-seq

For the mix condition 6 tibialis, 6 gastrocnemius, 6 soleus, 6 plantaris, and 6 EDL were pulled together. For the soleus condition, 24 soleus, for the tibialis condition 6 tibialis and for the quadriceps condition 6 quadriceps were pulled together. Muscles were dissected and pulled in cold PBS with 0,2U/μl RNase inhibitor. PBS was removed and muscles were minced in 1ml cold lysis buffer (10mM Tris-HCl pH7.5, 10mM NaCl, 3mM MgCl2, and 0.1% NonidetTM P40 in Nuclease-Free Water) with scissors. After 2min, 4ml of cold lysis buffer were added and muscles were lysed for 3 min at +4°C. 9ml of cold wash buffer (PBS, BSA 2% and 0.2U/μl RNase inhibitor from Roche) was added and the lysate was dounced with 10 strokes of loose pestle avoiding too much pressure and air bubbles. The homogenate was filtered with 70μm and 40μm cell strainers. Nuclei were pelleted by centrifugation for 5min at 500g at +4°C. Nuclei were washed to remove ambient RNAs in 1ml of cold wash buffer, transferred in a 1.5ml Eppendorf tube and centrifuged 5min at 500g at +4°C. The pellet was resuspended in 250 μl of wash buffer and stained during 15 min in the dark at +4°C with DAPI (10 μg/mL). Then the nuclei were washed with 1ml of wash buffer, centrifuged 5min at 500g at +4°C, resuspended in 300μl of wash buffer and filtered with 30μm cell strainers. Nuclei were then FACS sorted to exclude debris with a BD FACSAria III and the BD FACSDIVA software. More than 200 000 nuclei were collected in a tube containing 200 μl of wash buffer.

### Single-nucleus RNA-seq from skeletal muscle

After FACS sorting, nuclei were centrifuged 5min at 500g at +4°C, resuspended in 15μl of wash buffer and counted with a hemocytometer. The concentration of nuclei was then adjusted to 1000 nuclei/μl with wash buffer. We loaded around 4000 nuclei per condition into the 10X Chromium Chip. We used the Single Cell 3’ Reagent Kit v3 according to the manufacturer’s protocol. GEM-RT was performed in a thermal cycler: 53°C for 45 min, 85°C for 5 min. Post GEM-RT Cleanup using DynaBeads MyOne Silane Beads was followed by cDNA amplification (98°C for 3 min, cycled 12 x 98°C for 15 s, 67°C for 20s, 72°C for 1 min). After a cleanup with SPRIselect Reagent Kit and fragment size estimation with High SensitivityTM HS DNA kit runned on 2100 Bioanalyzer (Agilent), the libraries were constructed by performing the following steps: fragmentation, end-repair, A-tailing, SPRIselect cleanup, adaptor ligation, SPRIselect cleanup, sample index PCR, and SPRIselect size selection. The fragment size estimation of the resulting libraries was assessed with High SensitivityTM HS DNA kit runned on 2100 Bioanalyzer (Agilent) and quantified using the QubitTM dsDNA High Sensitivity HS assay (ThermoFisher Scientific). Libraries were then sequenced by pair with a HighOutput flowcel using an Illumina Nextseq 500 with the following mode:26base-pairs (bp) (10X Index + UMI), 8bp (i7 Index) and 57bp (Read 2).

### Single-nucleus RNA-seq analysis

A minimum of 50 000 reads per nucleus were sequenced and analyzed with Cell Ranger Single Cell Software Suite 3.0.2 by 10X Genomics. Raw base call files from the Nextseq 500 were demultiplexed with the cellranger mkfastq pipeline into library specific FASTQ files. The FASTQ files for each library were then processed independently with the cellranger count pipeline. This pipeline used STAR21 to align cDNA reads to the Mus musculus transcriptome (Sequence: GRCm38, Mouse reference provided by 10X.). Once aligned, barcodes associated with these reads –cell identifiers and Unique Molecular Identifiers (UMIs), underwent filtering and correction. Reads associated with retained barcodes were quantified and used to build a transcript count table. We re-runned this same pipeline with the pre-mRNA reference, build with cellranger mkgtf and cellranger mkref. Quality control on aligned and counted reads was performed keeping cells with > 500 reads, > 250 detected genes and less than 5 % mitochondrial genes. The subsequent visualizations, clustering and differential expression tests were performed in R (v 3.4.3) using Seurat36 (v3.0.2) (Stuart et al., 2019). We get 6477 nuclei from the mix of tibialis, EDL, gastrocnemius, plantaris and soleus, 1335 nuclei from quadriceps, 4838 nuclei from tibialis, 2517 nuclei from soleus. We detected the expression of approximately one thousand genes per nucleus in each of these muscles (Figure S1).

### Single-nucleus ATAC-seq from skeletal muscle

We used the 10X genomic nuclei Isolation for Single Cell ATAC Sequencing protocol (CG000169 | Rev B) with some changes. 12 quadriceps and 12 soleus were dissected and pulled in cold PBS. PBS was removed and muscles were minced 2 minutes in 1 ml of cold ATAC-lysis buffer (10mM Tris-HCl pH7.4, 10mM NaCl, 3mM MgCl2, 1% BSA and 0.1% Tween-20 in Nuclease-Free Water). 6ml of cold ATAC-lysis buffer were added and muscles were lysed on ice. After 3 minutes the lysate was dounced with 10 strokes of loose pestle avoiding too much pressure and air bubbles. After douncing, 8 ml of wash buffer were added and the homogenate was filtered with 70μm, 40μm and 20μm cell strainers. Nuclei were pelleted by centrifugation for 5min at 500g at +4°C. Next, we used the Chromium Single Cell ATAC kit according to the manufacturer’s protocol. Nuclei were resuspended in nuclei buffer from the kit, transposed 1 hour at 37°C. We loaded around 6000 nuclei into the 10X Chromium Chip. GEM incubation and amplification were performed in a thermal cycler: 72°C for 5 min, 98°C for 30 sec and 12 repeated cycles of 98°C for 10 sec, 59°C for 30 sec and 72°C for 1 min. Post GEM Cleanup using DynaBeads MyOne Silane Beads was followed by library construction (98°C for 45 sec, cycled 12 x 98°C for 20 sec, 67°C for 30 sec, 72°C for1 min). The libraries were constructed by adding sample index PCR, and SPRIselect size selection. The fragment size estimation of the resulting libraries was assessed with High SensitivityTM HS DNA kit runned on 2100 Bioanalyzer (Agilent) and quantified using the QubitTM dsDNA High Sensitivity HS assay (ThermoFisher Scientific). Libraries were then sequenced by pair with a HighOutput flowcel using an Illumina Nextseq 500.

### Single-nucleus ATAC-seq analysis

A minimum of 10 000 reads per nucleus were sequenced and analyzed with Cell Ranger Single Cell Software Suite 3.0.2 by 10X Genomics. Raw base call files from the Nextseq 500 were demultiplexed with the cellranger-atac mkfastq pipeline into library specific FASTQ files. The FASTQ files for each library were then processed independently with the cellranger count pipeline. This pipeline used STAR21 to align reads to the Mus musculus genome. Once aligned, barcodes associated with these reads –cell identifiers and Unique Molecular Identifiers (UMIs), underwent filtering and correction. The subsequent visualizations, clustering and differential expression tests were performed in R (v 3.4.3) using Seurat36 (v3.0.2) (Stuart et al., 2019), Signac (v0.2.4) (https://github.com/timoast/signac) and Chromvar (v1.1.1) (Schep et al., 2017). Quality control on aligned and counted reads was performed keeping cells with peak_region_fragments > 3000 reads and < 100000, pct reads in peaks > 15, blacklist ratio < 0.025, nucleosome_signal < 10 and TSS.enrichment > 2. We get 6037 nuclei in total and we detected 132 966 peaks (Figure. S1). The motif activity score was analyzed by running chromVAR (Figure. 3C-E).

### FISH with amplification (RNAscope) on isolated fibers

RNAscope^®^ Multiplex Fluorescent Assay V2 was used to visualize f*Myh* premRNAs and mRNAs, and *AldolaseA* and *Idh2* pre-mRNAs. Twenty different pairs of probes against the first intron of each f*Myh, AldolaseA* and *Idh2* pre-mRNAs were designed by ACDbio. Muscles were dissected and immediately fixed in 4% PFA at +4°C for 30 minutes. After fixation muscles were washed 3 times in PBS for 5 min. Myofibers were dissociated mechanically with small tweezers and fixed onto Superfrost plus slides (Thermo Fischer) coated with Cell-Tak (Corning) by dehydration at +55°C during 5 min. Slides were then proceeded according to the manufacturer’s protocol: ethanol dehydration, 10 min of H2O2 treatment and 30 min of protease IV treatment. After hybridization and revelation, the fibers were mounted under a glass coverslip with Prolong Gold Antifade Mountant (Thermofischer). Myofibers were imaged with a Leica DMI6000 confocal microscope composed by an Okogawa CSU-X1M1 spinning disk and a CoolSnap HQ2 Photometrics camera. Images were analyzed with Fiji Cell counter program.

### Immunohistochemistry

For immunostaining against MYH4, MYH2, MYH7 and Laminin, adult legs without fixation and without skin were embedded with TissuTEK OCT and directly frozen in cold isopentane cooled in liquid nitrogen. Muscles were conserved at −80°C and cut with Leica cryostat 3050s with a thickness of 10μm. Cryostat sections were washed 3 times 5 minutes with PBS and then incubated with blocking solution (PBS and 10% goat serum) 30 minutes at room temperature. Sections were incubated overnight with primary antibody solution at +4°C, then washed 3 times for 5 minutes with PBS and incubated with secondary antibody solution 1 hour at room temperature. Sections were further washed 3 times 5 minutes and mounted with Mowiol solution and a glass coverslip. Images were collected with an Olympus BX63F microscope and a Hamamatsu ORCA-Flash 4.0 camera. Images were analyzed with ImageJ program. The references of the antibodies used are listed in Table 1.

### Statistical analysis

The graphs represent mean values ± SEM. Significant differences between mean values were evaluated using two-way ANOVA with Graphpad 6 software and student t test for Figure Sup 9C.

### GEO data accession number

Single nucleus RNA-seq and Single nucleus ATAC-seq data will be deposited in the NCBI Gene Expression Omnibus database (https://www.ncbi.nlm.nih.gov/geo/).

