## Supplemental for "Single-nucleus RNA-seq and FISH reveal coordinated transcriptional activity in mammalian myofibers"

FIGURE_S1_DOS_SANTOS_ET_AL_2020

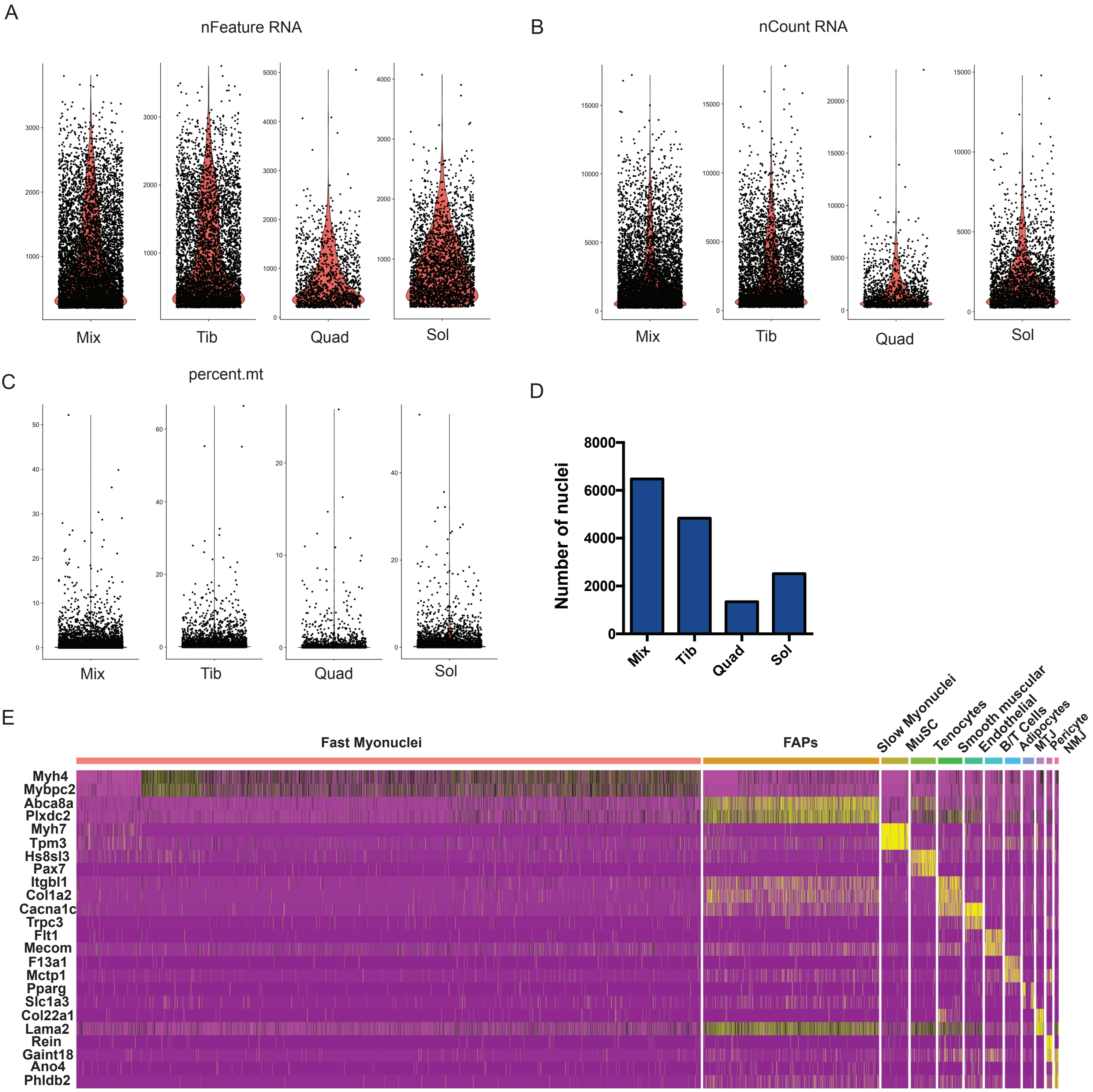


**Figure S1. Quality controls of the snRNA-seq experiments from Figure 1.**

(A) Number of detected genes per sample using exonic and intronic sequences. (B) Number of reads per sample using exonic and intronic sequences. (C) Percentage of mitochondrial genes detected per sample. (D) Number of nuclei per sample after selecting nuclei that have unique feature counts between 2,500 and 200 and less than 5% mitochondrial counts. For each panel: Mix: mix of tibialis, EDL, gastrocnemius, plantaris and soleus; Tib: Tibialis; Quad: Quadriceps; Sol: Soleus. (E) Heatmap of the top two genes preferentially expressed (yellow) in each population of nuclei.

FIGURE_S2_DOS_SANTOS_ET_AL_2020

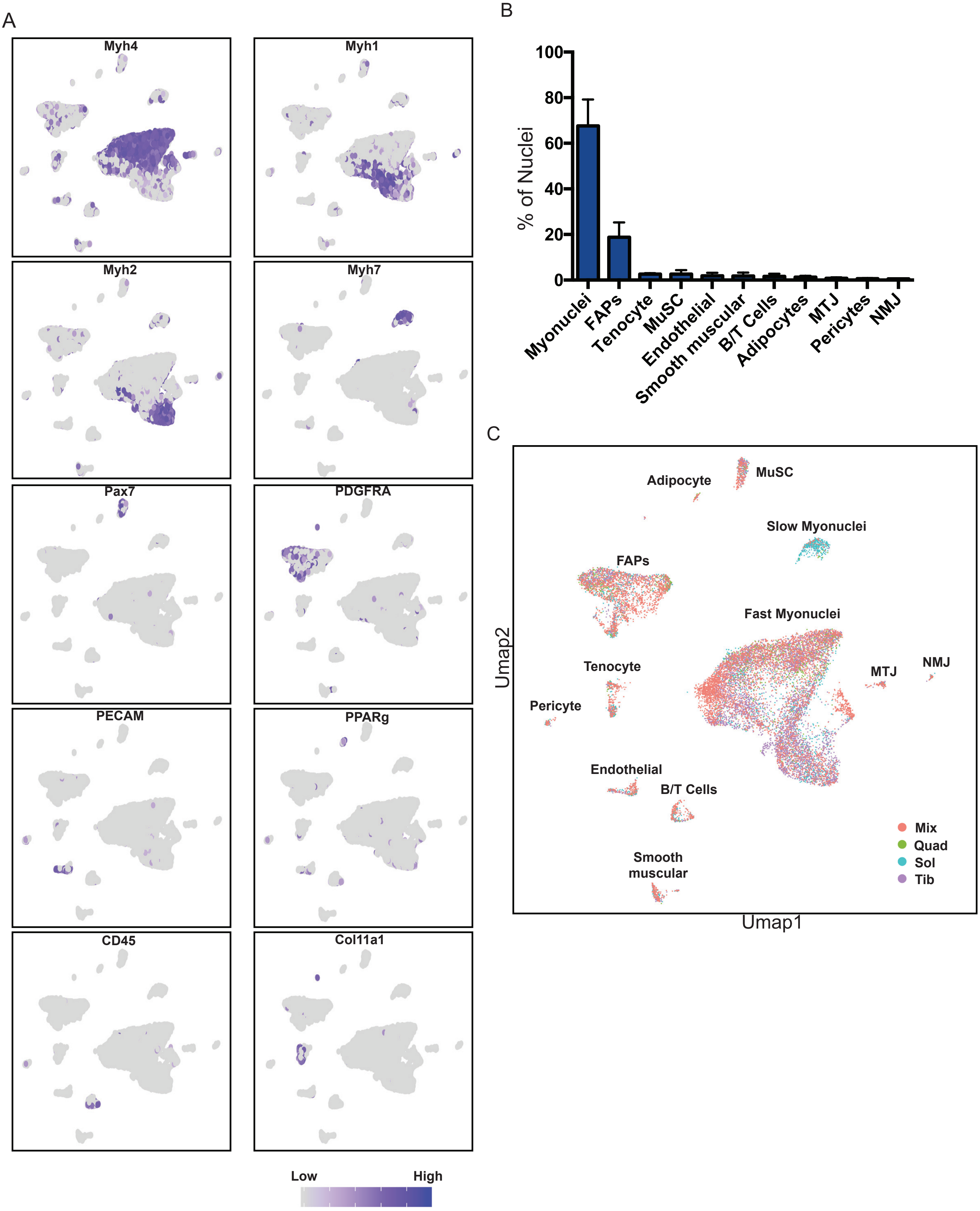

**Figure S2. Origin and identity of the nuclei populations defined in Figure 1**

(A) Same Umap diagram than in Figure 1B, showing the expression of several markers used to identify the different cell types populations. The intensity of the blue color depends on the level of expression of the gene. (B) Quantification of the percentage of cell types found by snRNA-seq in the distinct studied muscles. (C) Same Umap diagram than in Figure 1B showing the origin of each nucleus. MuSC: Skeletal muscle stem cells; FAPs: Fibro-adipogenic progenitors; MTJ: myotendinous junction; NMJ: neuromuscular junction.

FIGURE_S3_DOS_SANTOS_ET_AL_2020

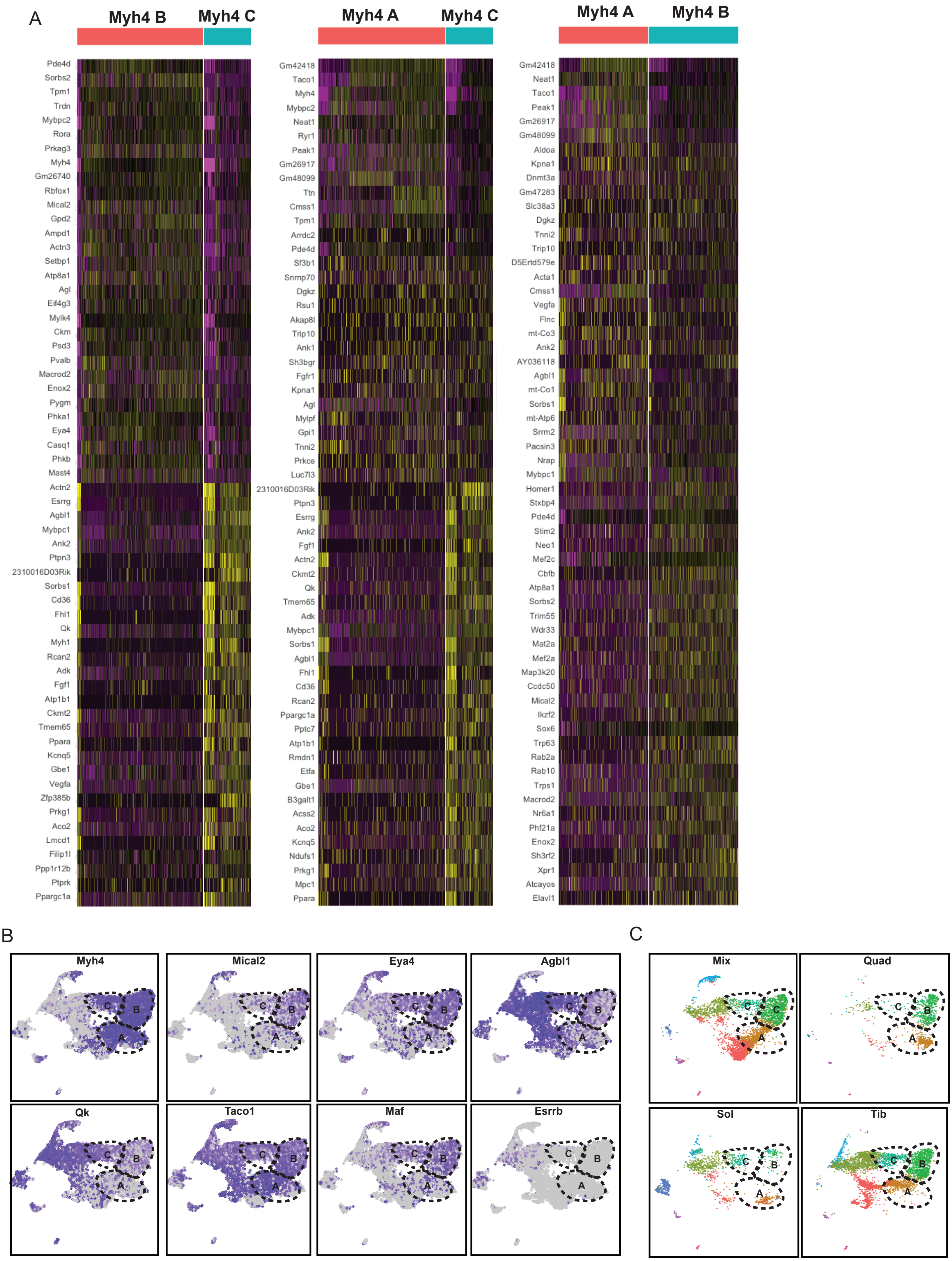


**Figure S3. The different populations of *Myh4*+ myonuclei.**

(A) Heatmap of genes upregulated (yellow) and downregulated (violet) in the different clusters of *Myh4* positive nuclei. (B) Same Umap diagram than in Figure 1C showing the expression of several genes differentially expressed in the 3 sub-clusters of *Myh4* myonuclei. The intensity of the violet color depends on the gene expression level. (C) Same Umap diagram than Figure 1C showing the origin of each nucleus.

FIGURE_S4_DOS_SANTOS_ET_AL_2020

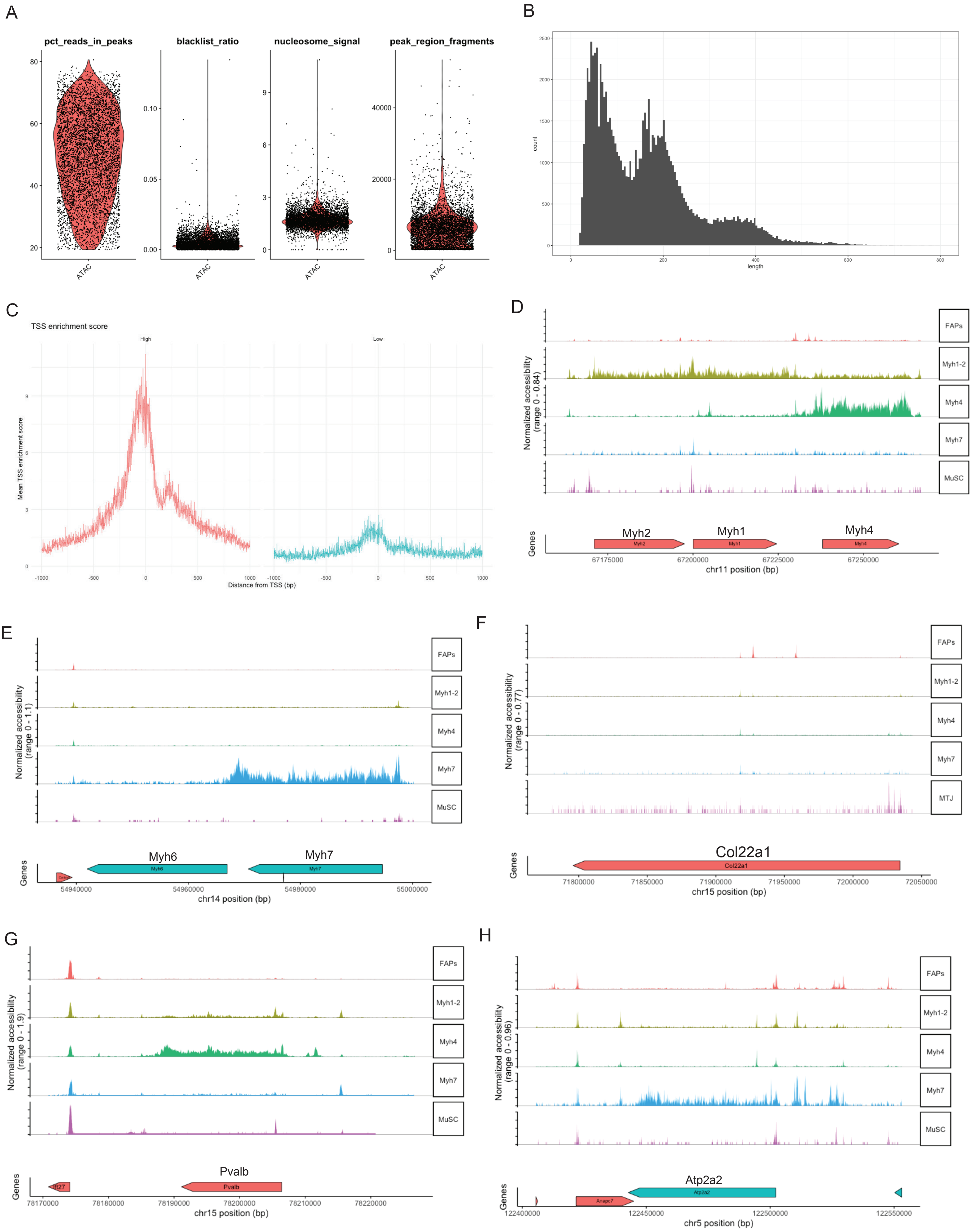

**Figure S4. Quality controls of the snATAC-seq experiments from Figure 3.**

(A) Violin plot from snATAC-seq experiments from Figure 3, showing the percentage of reads in peaks, the ratio of reads in blacklist region versus peaks region, the nucleosome signal and the peak region fragments. (B) Histogram showing the fragment length periodicity for all the nuclei from snATAC-seq experiments. (C) Histogram showing the enrichment of the peaks from snATAC-seq at transcriptional start sites (TSSs). (D) Chromatin accessibility in the f*Myh* locus in the different types of nuclei present in skeletal muscles. (E) Same as (D) for the slow *Myh7* locus. (F) Same as (D) for the *Col22a1* locus. (G) Same as (D) for the *Pvalb* locus. (H) Same as (D) for the *Atp2a2* locus.

FIGURE_S5_DOS_SANTOS_ET_AL_2020

A

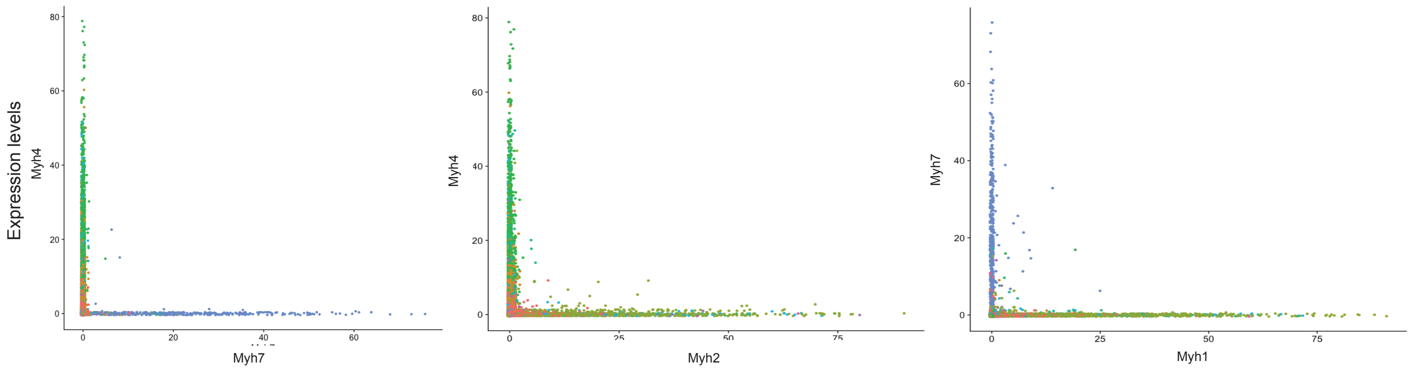


**Figure S5. The majority of myonuclei express only one isoform of *Myh*.**

(A) Analysis of *Myh* isoforms expression in myonuclei from snRNA-seq data. Each dot corresponds to a myonucleus and the x- and y-axis corresponds to the indicated *Myh* expression level.

FIGURE_S6_DOS_SANTOS_ET_AL_2020

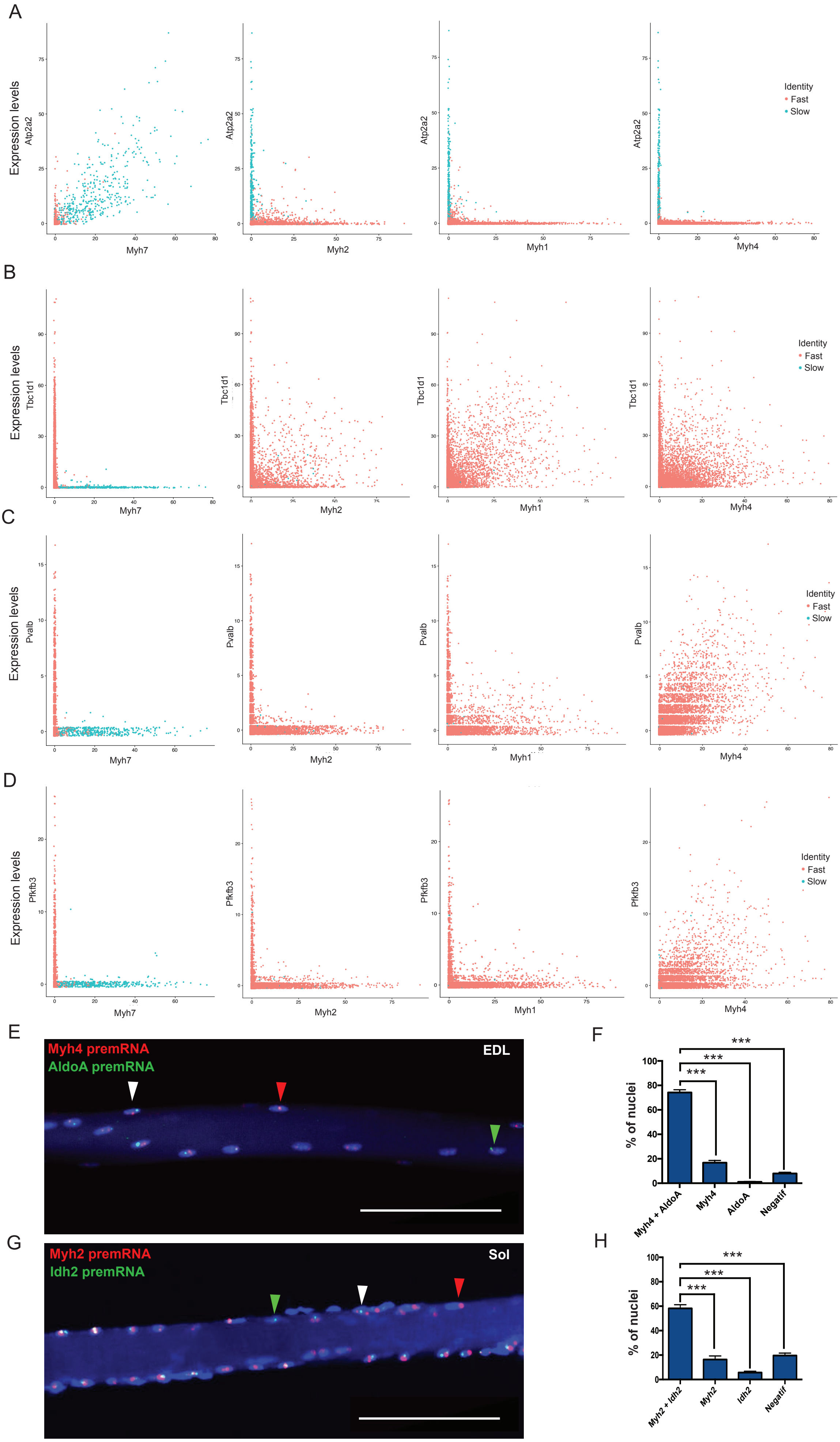

**Figure S6. Sarcomeric and metabolic genes are co-expressed in myonuclei.**

(A) Analysis of *Atp2a2* and *Myh* isoform expression in myonuclei from snRNA-seq data from Figure 1A. Each dot corresponds to one myonucleus and the axis correspond to the gene expression level. *Atp2a2* is expressed only in most *Myh7* and in few *Myh2* nuclei. (B) Same as (A) with *Tbc1d1* expression. *Tbc1d1* is expressed only in fast and not in slow nuclei. (C) Same as (A) with *Pvalb* expression. *Pvalb* is expressed only in *Myh1* and *Myh4* nuclei. (D) Same as (A) with *Pfkfb3* expression. *Pfkfb3* is expressed preferentialy in *Myh4* nuclei. (E) RNAscope on isolated fibers from EDL showing the localization of *Myh4* (red) and *AldoA* (green) pre-mRNAs. Most myonuclei expressed at the same time *Myh4* and *AldoA* (white arrowhead). But some nuclei expressed *Myh4* without *AldoA* (red arrowhead), and some nuclei are negative for both genes. (F) Percentage of the nuclei co-expressing *Myh4* and *AldoA* or expressing only *Myh4* or *AldoA* and negative nuclei in EDL fibers. (G) RNAscope on isolated fibers from soleus showing the localization of *Myh2* (red) and *Idh2* (green) pre-mRNAs. Like in (E) most myonuclei expressed at the same time *Myh2* and *Idh2* (white arrowhead). But some nuclei expressed *Myh2* without *Idh2* (red arrowhead) and other nuclei are negative for both genes. (H) Percentage of the nuclei co-expressing *Myh2* and *Idh2* or expressing only *Myh2* or *Idh2* and negative nuclei in soleus fibers. For E-H scale bar: 100µm. For F-H. The graphs show data pooled from three animals and fifteen fibers per animal. Numerical data are presented as mean ± s.e.m. *P < 0.05, **P < 0.01, ***P < 0.001.

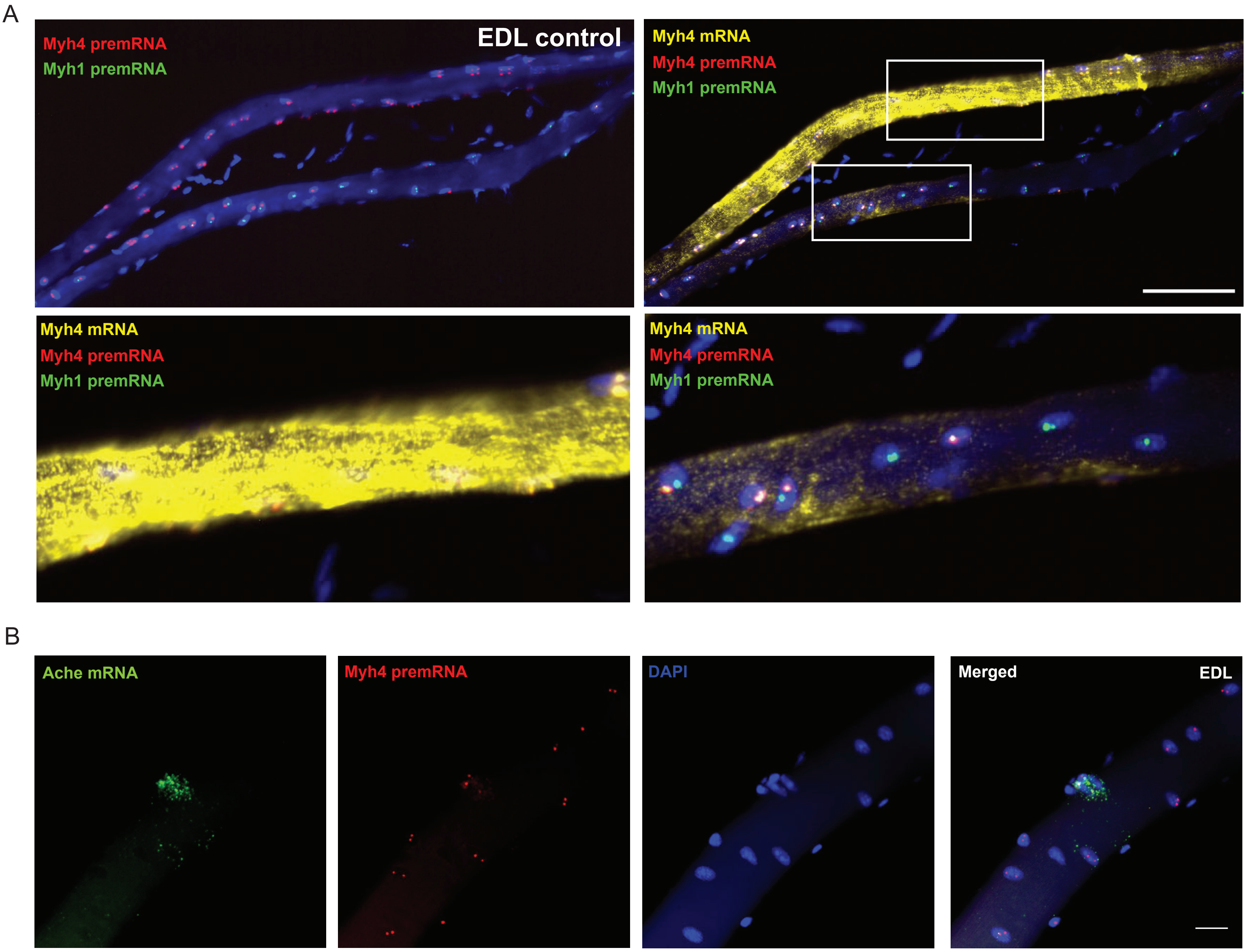


**Figure S7. Transcriptional variability of *Myh* expression all along myofibers** (A) Up: RNAscope on isolated fibers from EDL showing the localization of *Myh4* (red), *Myh1* (green) pre-mRNAs and *Myh4* (yellow) mRNAs in pure and hybrid fibers. Down left: the coordinated fiber shows a homogeneous accumulation of *Myh4* mRNA (in yellow) all along the myofiber. Down right: the hybrid fiber shows an accumulation of *Myh4* mRNA around *Myh4* positive nuclei (arrowhead) but not around *Myh1* positive nuclei. (B) RNAscope showing the expression of *Ache* mRNA and *Myh4* pre-mRNA in NMJ nuclei (arrowhead). For A scale bar: 100µm. For B scale bar: 20µm.

FIGURE_S8_DOS_SANTOS_ET_AL_2020

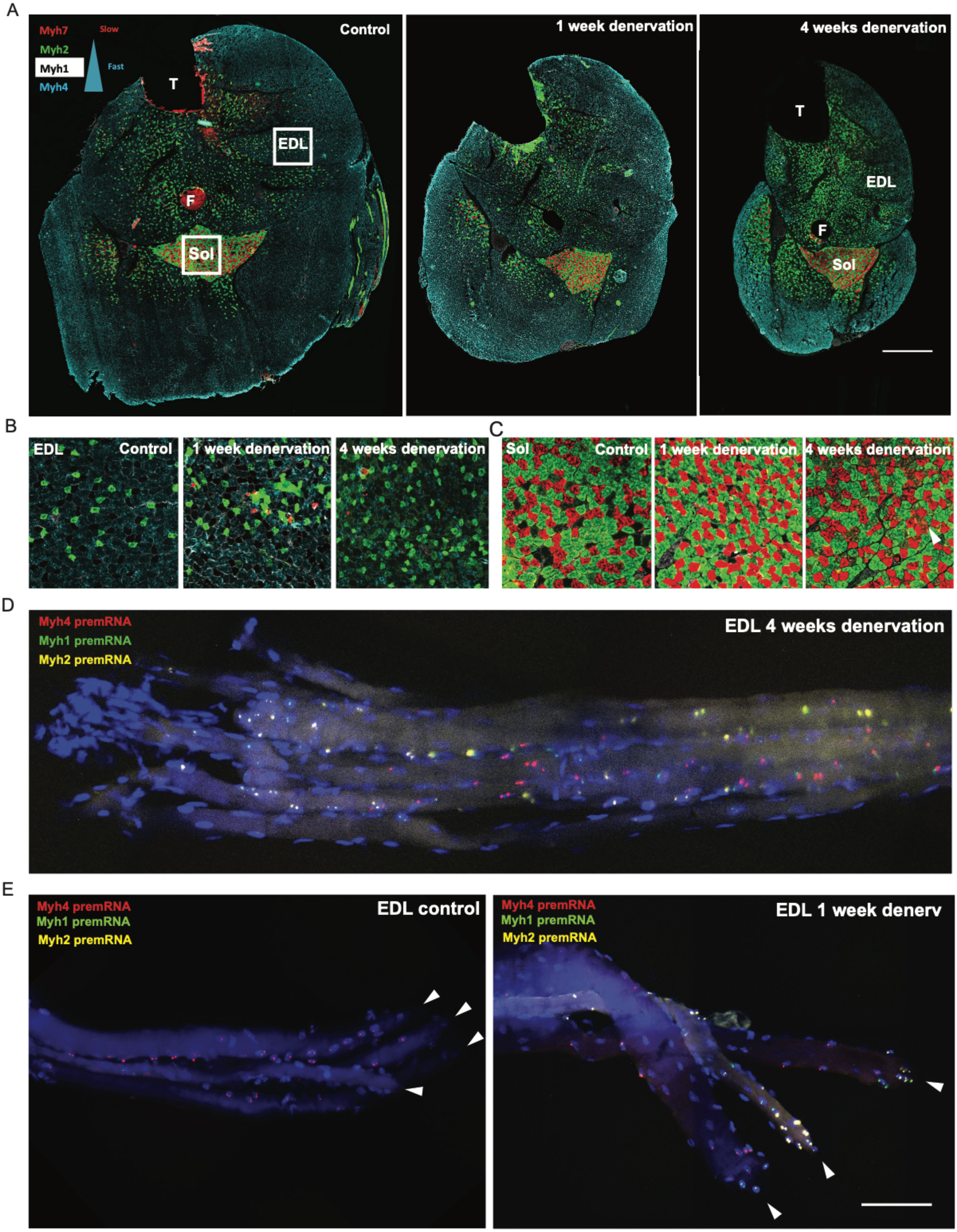

**Figure S8. *Myh* expression during denervation.**

(A) Immunostaining of MYH proteins in a section of the leg of an adult mouse after 1 week (center) and 4 weeks of denervation (right) and control (left). During denervation, fast MYH4 fibers located on the periphery of the leg show preferential atrophy compared to the slower fibers in the center of the leg. Fibers from all muscles become slower (fast to slow transition) : muscles have less MYH4 (blue) fibers and more MYH1 (black), MYH2 (green) and MYH7 (red) fibers. T: Tibia; F: Fibula; Sol: Soleus; EDL: Extensor digitalis longus. (B) Same as (A). Zoom in EDL muscle. (C) Same as (A). Zoom in soleus muscle. (D) RNAscope on isolated fibers from 4 weeks denervated EDL, showing the localization of *Myh4* (red), *Myh1* (green) and *Myh2* (yellow) pre- mRNAs. After 4 weeks of denervation, fibers are still de-coordinated. (E) RNAscope on isolated fibers from control and 1 week denervated EDL, showing the localization of *Myh4* (red), *Myh1* (green) and *Myh2* (yellow) pre-mRNAs. Left: Nuclei from the myotendinous junctions of control EDL fibers express the same *Myh* isoform than other nuclei (arrowhead). Right: After 1 week of denervation, MTJ nuclei (arrowhead) express different isoforms of *Myh* than the other myonuclei, and this expression seems more important than in the other nuclei.

FIGURE_S9_DOS_SANTOS_ET_AL_2020

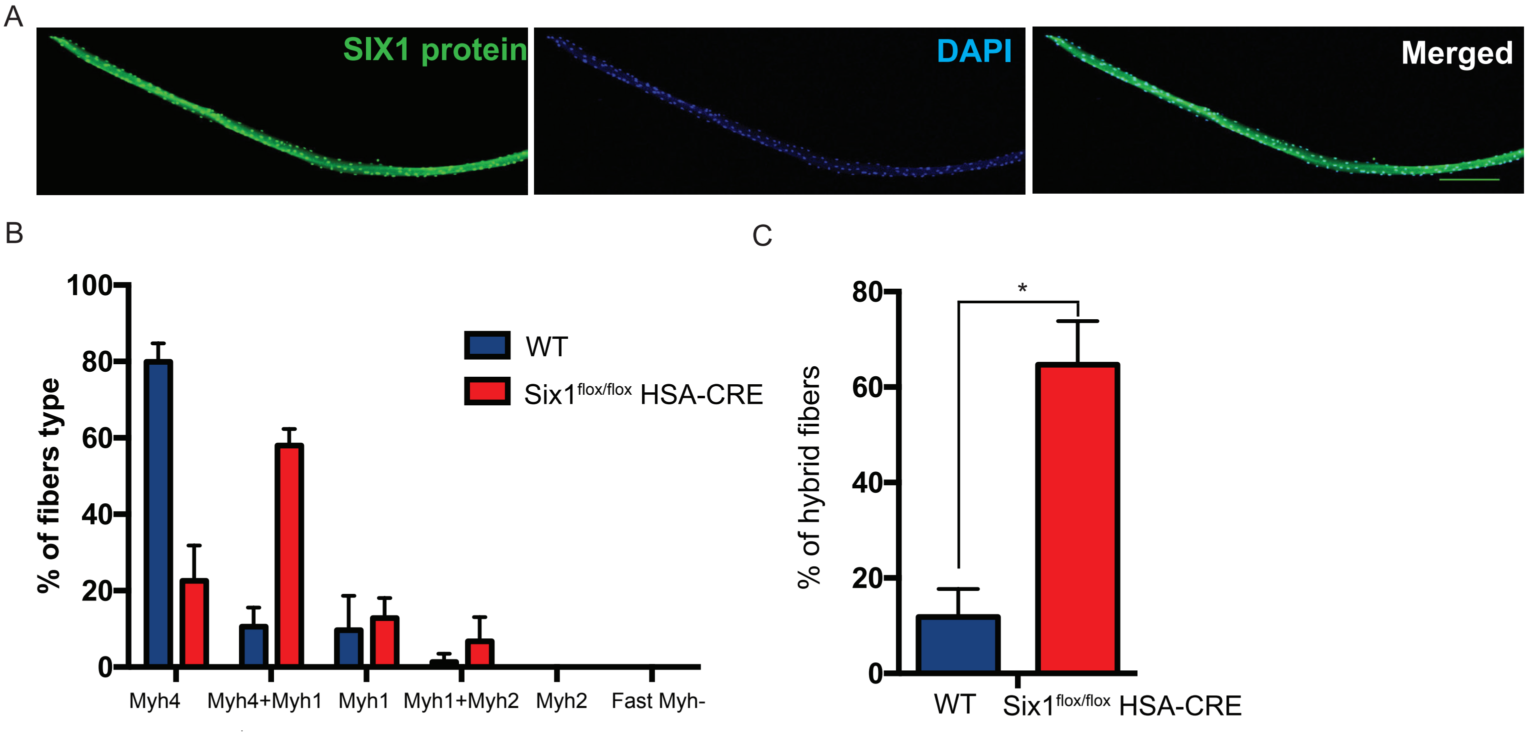


**Figure S9. Coordination of *Myh* expression throughout the fiber is lost in *Six1* mutant myofibers.**

(A) SIX1 accumulates in all myonuclei of adult EDL fibers as revealed by immunohistochemistry with SIX1 specific antibodies. (B) Percentage of the different types of EDL fibers observed in WT and *Six1*$^{flox/flox}$ *;HSA-CRE* mice by RNAscope experiments. (C) Percentage of hybrid EDL fibers (de-coordinated) in WT and *Six1*$^{flox/flox}$ *;HSA-CRE* mice. The graphs show data pooled from three animals and more than fifty fibers in total. Numerical data are presented as mean ± s.e.m. *P < 0.05, **P < 0.01, ***P < 0.001.

FIGURE_S10_DOS_SANTOS_ET_AL_2020

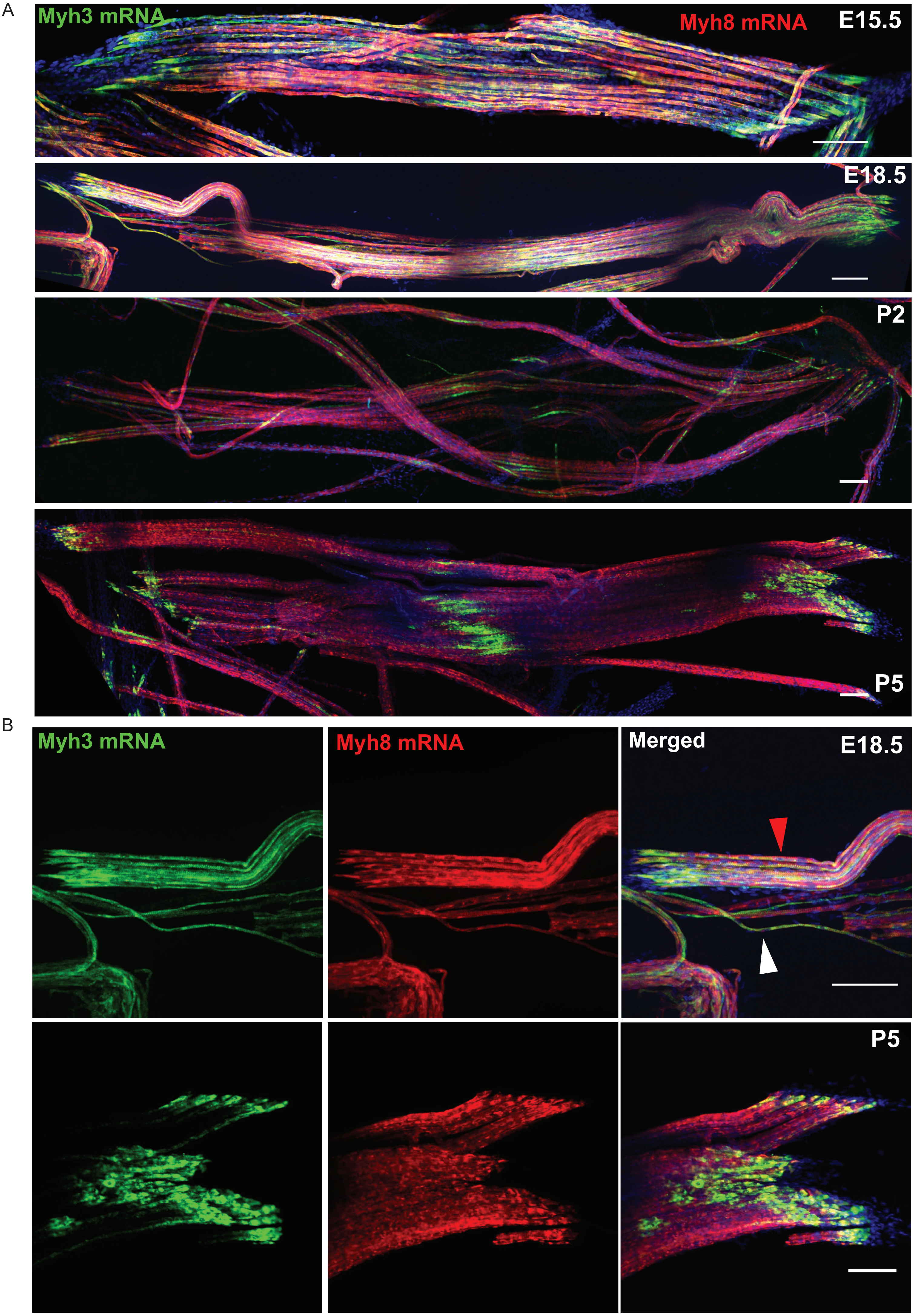

**Figure S10. *Myh3* expression during development is regionalized in MTJ nuclei in contrast to *Myh8* expression.**

(A) RNAscope against *Myh3* (green) and *Myh8* (red) mRNA on isolated fibers from forelimbs at E15.5, E18.5, P2 and P5. At E15,5 embryonic myofibers express *Myh3* and *Myh8*. The myotendinous junction areas show an accumulation of *Myh3* mRNA. At E18,5, fetal myofibers (small fibers) expressed more *Myh3* than embryonic fibers (big fibers). (B) Same as (A). Zoom in MTJ areas at E18,5, and P5. *Myh3* (green) accumulated in MTJ areas in contrast to *Myh8* (red). White arrowhead shows a primary myofiber and the red arrowhead a secondary myofiber.

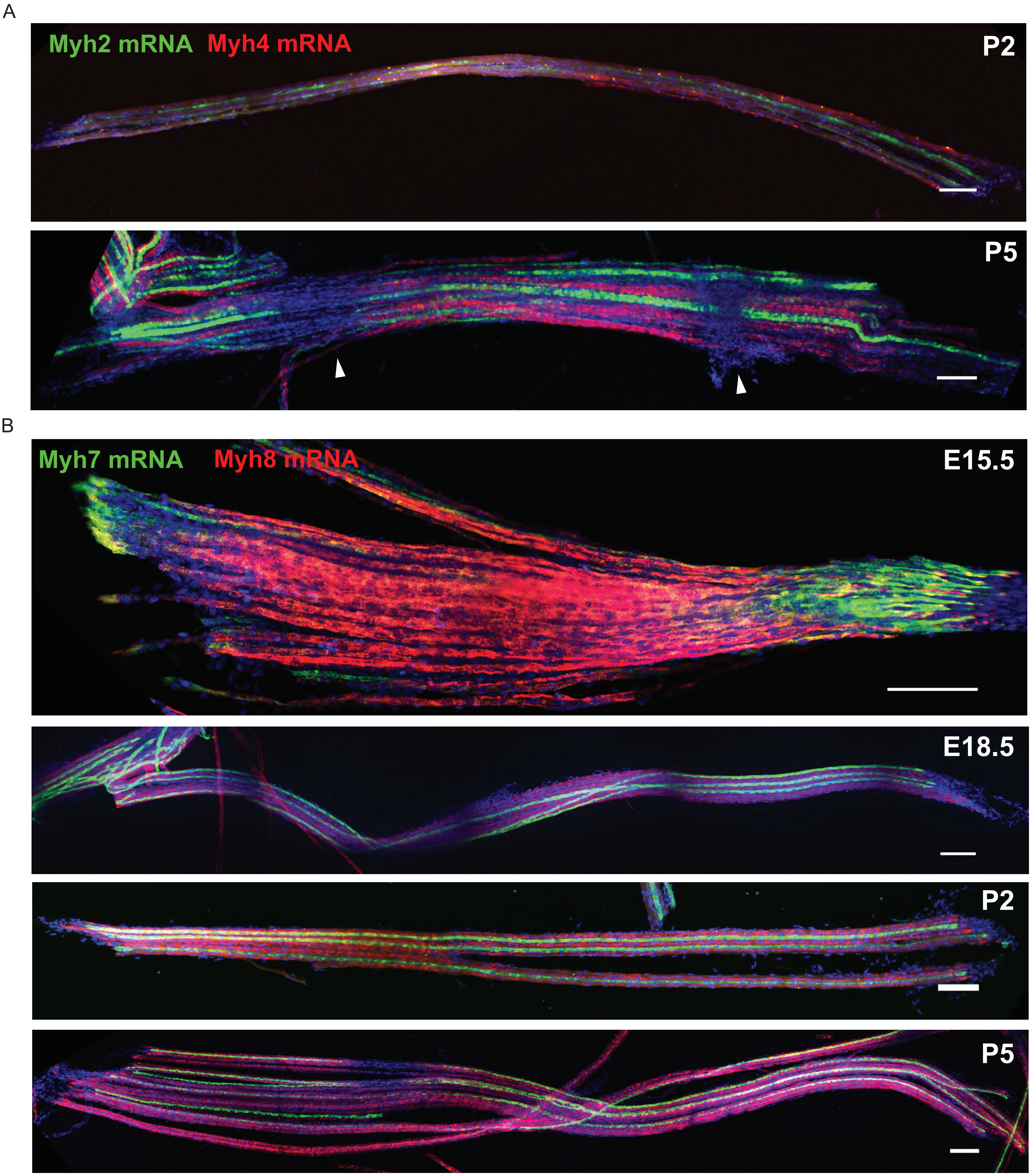


**Figure S11.** ***Myh7, Myh2 and Myh4* expression during development.**

(A) RNAscope against *Myh4* (red) and *Myh2* (green) mRNAs on isolated fibers at P2 and P5. The adult fast *Myh2* and *Myh4* mRNAs start to be detectable in some myofiber regions at P2.

The expression is strongly increased at P5 and is localized homogeneously along the myofiber. We did not detect hybrid *Myh4* and *Myh2* myofibers. Note that some parts of the myofibers at P5 do not present *Myh2* nor *Myh4* mRNA accumulation (arrowhead). (B) RNAscope against *Myh7* (green) and *Myh8* (red) mRNAs on isolated fibers at E15.5, E18.5, P2 and P5. *Myh7* mRNAs accumulate in MTJ areas of all myofibers at E15.5. After E18.5, *Myh7* mRNAs are detected homogeneously in slow myofibers and no more accumulating in MTJ areas.
